# VegVault dataset: linking global paleo-, and neo-vegetation data with functional traits and abiotic drivers

**DOI:** 10.1101/2025.01.31.635213

**Authors:** Ondřej Mottl, Franka Gaiser, Irena Šímová, Suzette G. A. Flantua

## Abstract

Understanding the dynamics and persistence of biodiversity patterns over short (contemporary) and long (thousands of years) time scales is crucial for predicting ecosystem changes under global climate and land-use changes. A key challenge is integrating currently scattered ecological data to assess complex vegetation dynamics over time. Here, we present VegVault, an interdisciplinary SQLite database that uniquely integrates paleo- and neo-ecological plot-based vegetation data on a global and millennial scale, directly linking them with functional traits, soil, and climate information. VegVault currently comprises data from *BIEN*, *sPlotOpen*, *TRY*, *Neotoma*, *CHELSA*, and *WoSIS*, providing a comprehensive and ready-to-use resource for researchers across various fields to address questions about past and contemporary biodiversity patterns and their abiotic drivers. To further support the usability of the data, VegVault is complemented by the {*vaultkeepr*} R package, enabling streamlined data access, extraction, and manipulation. This study introduces the structure, content, and diverse applications of VegVault, emphasizing its potential role in advancing ecological research to improve predictions of biodiversity responses to global climate change.

## Background & Summary

Biodiversity and ecosystem dynamics are currently profoundly influenced by global climate change, creating an urgent need for robust predictive models capable of anticipating these impacts. Such models are crucial in guiding conservation efforts during this global crisis. However, many existing models rely on contemporary (present day) ecological data alone, which limits their ability to capture the long-term responses of species and ecosystems to environmental change. Incorporating paleoecological data—“ecology of the past” as opposed to neo-ecological, or “ecology of the present”—offers a vital temporal perspective that can enhance our understanding of biodiversity dynamics under shifting climate and land-use^1–3^.

In recent years, an increasing number of large-scale ecological datasets have become available, providing unprecedented opportunities to investigate ecosystem dynamics across spatial and temporal scales. These datasets encompass contemporary vegetation data (e.g., *Global Biodiversity Information Facility*^4^ (‘*GBIF*’ hereafter), *sPlot*^5^, *Botanical Information and Ecology Network*^6^ (‘*BIEN*’ hereafter), paleo-vegetation data (e.g., *Neotoma Paleoecology Database*^7^ (‘*Neotoma*’ hereafter)), plant functional traits (e.g., *TRY plant trait database*^8^ (‘*TRY*’ hereafter)), and abiotic drivers (e.g., *CHELSA*^9^, *WorldClim*^10^), which are all critical for understanding the complex interactions that shape ecosystems. Yet, despite the growing availability of such datasets, efforts to integrate these diverse sources into a unified, open-access framework remain limited. An integrated database, built with reproducible workflows, is a crucial step toward improving predictive models that estimate biodiversity dynamics under changing climatic conditions. Addressing this gap requires interdisciplinary solutions^11,12^ and a commitment to compiling, processing, and linking data in a systematic and transparent manner. Currently, no comprehensive framework exists to link global vegetation data across temporal scales with functional traits and abiotic drivers.

There have been previous attempts to integrate vegetation data across extensive time scales and link them with other data types. For example, *BioDeepTime*^13^ compiles both contemporary and fossil biodiversity time series. However, it lacks coverage of contemporary vegetation plots and, more importantly, omits direct links to functional traits and abiotic data. GIFT^14^ includes vegetation biodiversity linked with functional traits but only holds taxonomic lists for specific regions, without data on relative abundances. Additionally, the latter database lacks the long-term dimension of vegetation dynamics over time, derived from the currently existing large databases containing paleoecological records, such as the Neotoma database.

**VegVault** addresses these limitations by offering a unique resource that compiles extensive paleo- and neo-ecological vegetation data into a single, accessible SQLite database, accompanied by functional traits and abiotic information (i.e., climate and soil). Its design bridges paleo- and contemporary data, enabling the study of long-term processes, and linking them to today’s patterns. Thereby, VegVault offers the possibility to specifically analyse the mechanisms of ecosystem responses to environmental change, which enhances our ability to study and ultimately predict future biodiversity trajectories. See **Table 1** for the comparative table that summarises the strengths and weaknesses of existing databases versus VegVault. Specifically, VegVault advances analyses across spatial, temporal, and taxonomic scales in several ways:

- **Contemporary vegetation data**: VegVault incorporates plot-based data from *BIEN*^6^ and *sPlotOpen*^15^ (open-access version of sPlot), including essential “true negative” information about species’ absence. These datasets are pre-cleaned for taxonomic and georeferencing errors, ensuring high reliability for subsequent analyses. The focus on plot-based data rather than occurrence data (e.g., *GBIF*) provides a closer match to paleoecological data formats, facilitating robust comparisons across temporal scales.
- **Paleo-vegetation data**: VegVault employs standardised fossil pollen data processed using the *FOSSILPOL* workflow^16^ to integrate records globally from Neotoma. This workflow ensures quality control and creates “ready-for-analysis” data through a reproducible and detailed process for handling large fossil pollen dataset compilations (see **Methods** for details).
- **Interdisciplinary integration**: VegVault combines past and present vegetation data with functional traits and abiotic information to support comprehensive ecological analyses. The abiotic information includes soil data from the *WoSIS Soil Profile Database*^17^ by *World Soil Information Service* (‘*WoSIS*’ hereafter), and global temperature and precipitation datasets from *CHELSA* and *CHELSA-TRACE21*^18,19^, providing essential environmental context. Functional trait data from the *TRY* database and *BIEN* enrich the database by linking species characteristics to ecological roles and responses. This allows researchers to move beyond taxonomic diversity and explore functional dimensions, offering deeper insights into ecosystem processes.
- **Taxonomic classification**: VegVault ensures consistent and standardised taxonomic information by aligning all taxa present in vegetation data—both paleo and contemporary— with the GBIF backbone taxonomy^20^. This process, implemented via the {*taxospace*} R package^21^, automates taxonomic classification by matching names to an existing taxonomy based on similarity (e.g., “Ulmus Type” will be classified as genus *Ulmus*). While original taxonomic names are preserved within the VegVault, users can access data at varying taxonomic resolutions (e.g., species, genus, family; if available) to suit specific analytical needs. The same standardisation approach is applied to vegetation and trait data, facilitating seamless integration between datasets. For instance, enabling the calculation of genus-level average trait values, when species-level identification is unavailable.
- **Accessibility**: VegVault is designed with user accessibility in mind. The SQLite database is supported by the {*vaultkeepr*} R package^22^, which simplifies and streamlines the process of data extraction for project-specific needs. It ensures that researchers can efficiently retrieve and subsequently manipulate data in a reproducible and transparent manner.

**Table 1:**
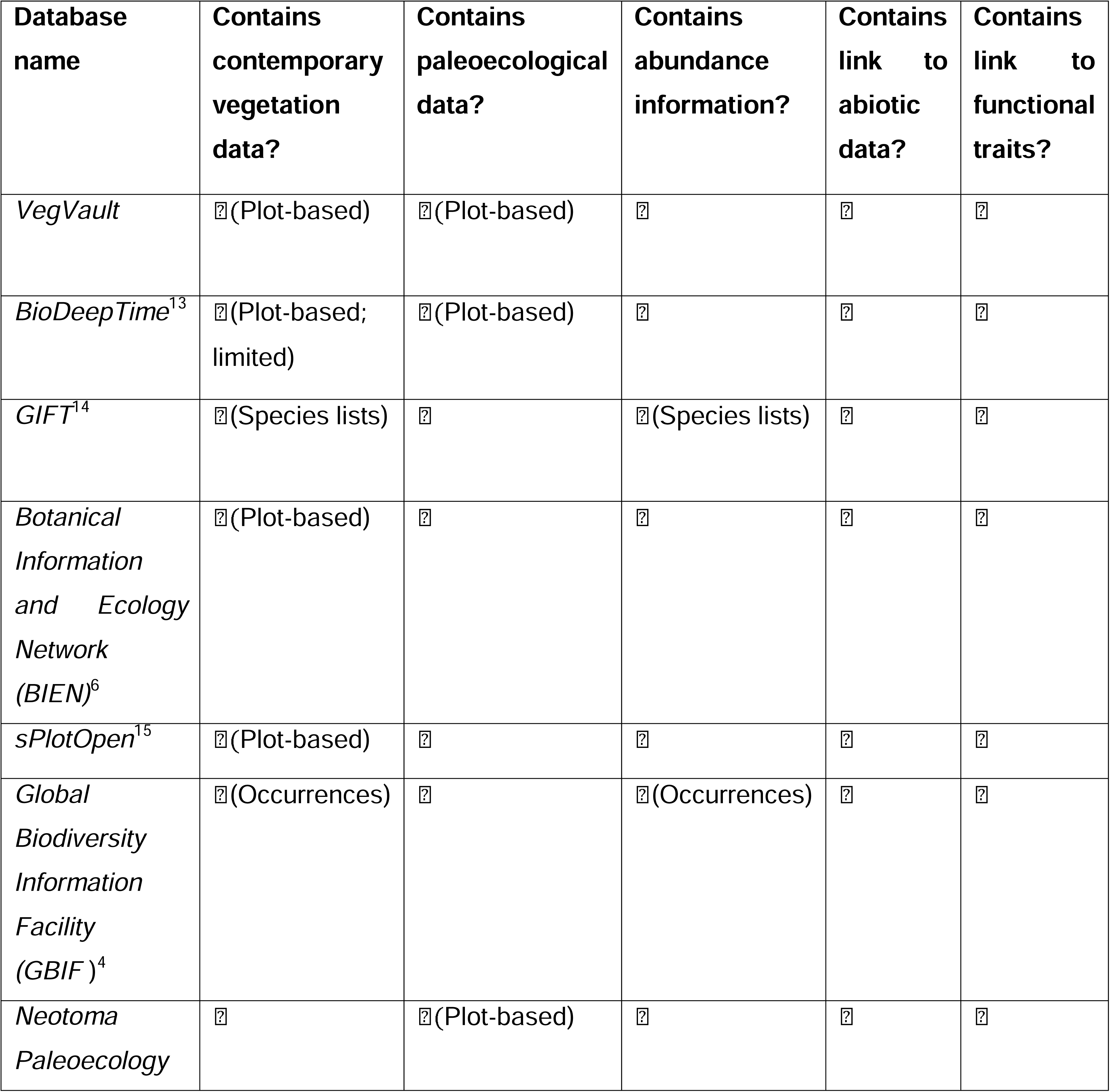
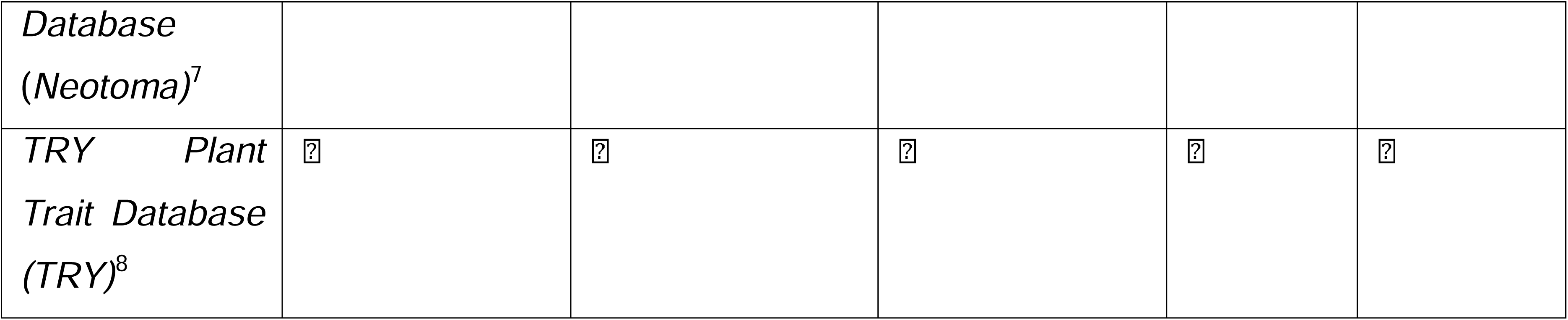
Overview comparison of selected databases and VegVault to highlight the latter’s novel contributions. Note that this comparison does not intend to be a comprehensive overview of all ecological databases. We specify the presence of “Plot-based” vegetation data and Species lists to flag the differences between stored data. The “Plot-based” databases contain a full survey of the vegetation at specific locations and, therefore, contain true negatives. Moreover, “Plot-based” databases contain some expression of species abundance.

**Table 2:**
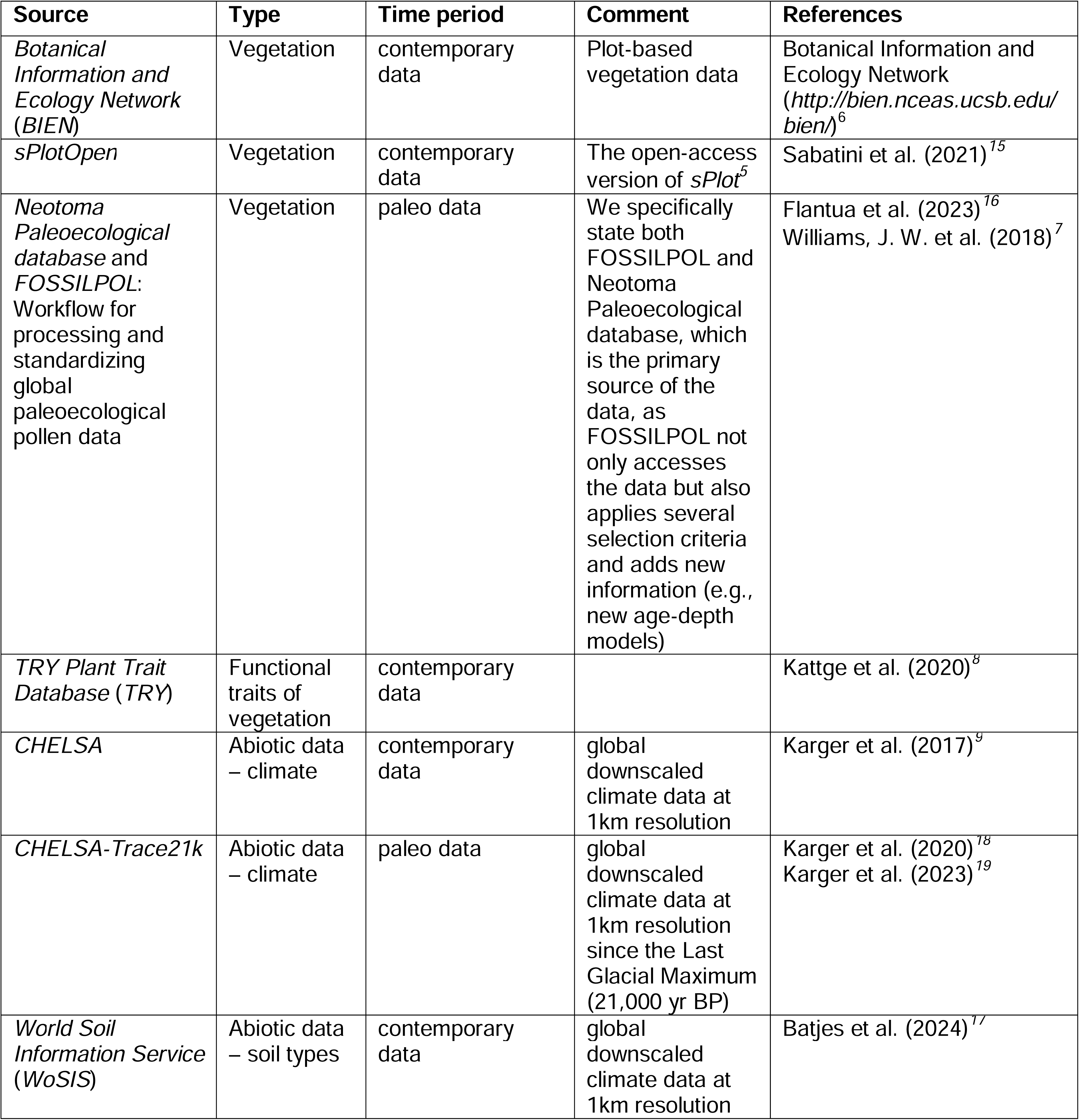
Main sources of data presented in VegVault v1.0.0.

The comprehensive nature of VegVault enables diverse applications across scientific disciplines. Its integration of fossil pollen data and functional traits allows researchers to investigate long-term vegetation dynamics and their responses to past climate variations. Such integration provides critical context for past ecological trends^e.g.,23,24^, including trait information, which is currently underrepresented in paleoecological analyses of vegetation dynamics^25^. While there are now studies showing the relationship between functional traits and climate for global contemporary data^26^, VegVault allows testing the universality of such relationships across temporal scales. In addition, VegVault provides the needed data for advanced modelling approaches, such as comparing species distributions across temporal scales^e.g.,27^, and analysing biodiversity patterns using Joint Species Distribution Models^28^. VegVault also facilitates studies of functional ecology and biogeography by enabling assessments of shifts in ecosystem functioning and climatic drivers over time, and by identifying the taxonomic and functional novelty of communities undergoing climate-driven changes^e.g.,29,30^. Furthermore, the detailed climate data (past and present) linked with vegetation records provide a robust foundation for examining the impacts of past and present climate change on ecosystems^e.g.,31^. These capabilities make VegVault a valuable interdisciplinary resource for predicting biodiversity patterns and informing conservation strategies and policy decisions aimed at mitigating climate change effects. While this is not an exhaustive list of potential applications, VegVault offers a rich and accessible resource that can significantly advance ecological research and global biodiversity conservation.

## Methods

### VegVault workflow and reproducibility

VegVault integrates data from multiple well-established primary sources to provide a comprehensive view of vegetation dynamics. The current sources of data are summarised in **Table 1**.

The data processing and integration into VegVault follow a meticulous workflow to ensure consistency and reliability. Each data source is first accessed and processed individually using a specific pipeline. See **Figure 1** for the workflow process of accessing and processing all data sources (1 to 6 in **Figure 1**).

**Figure 1.**
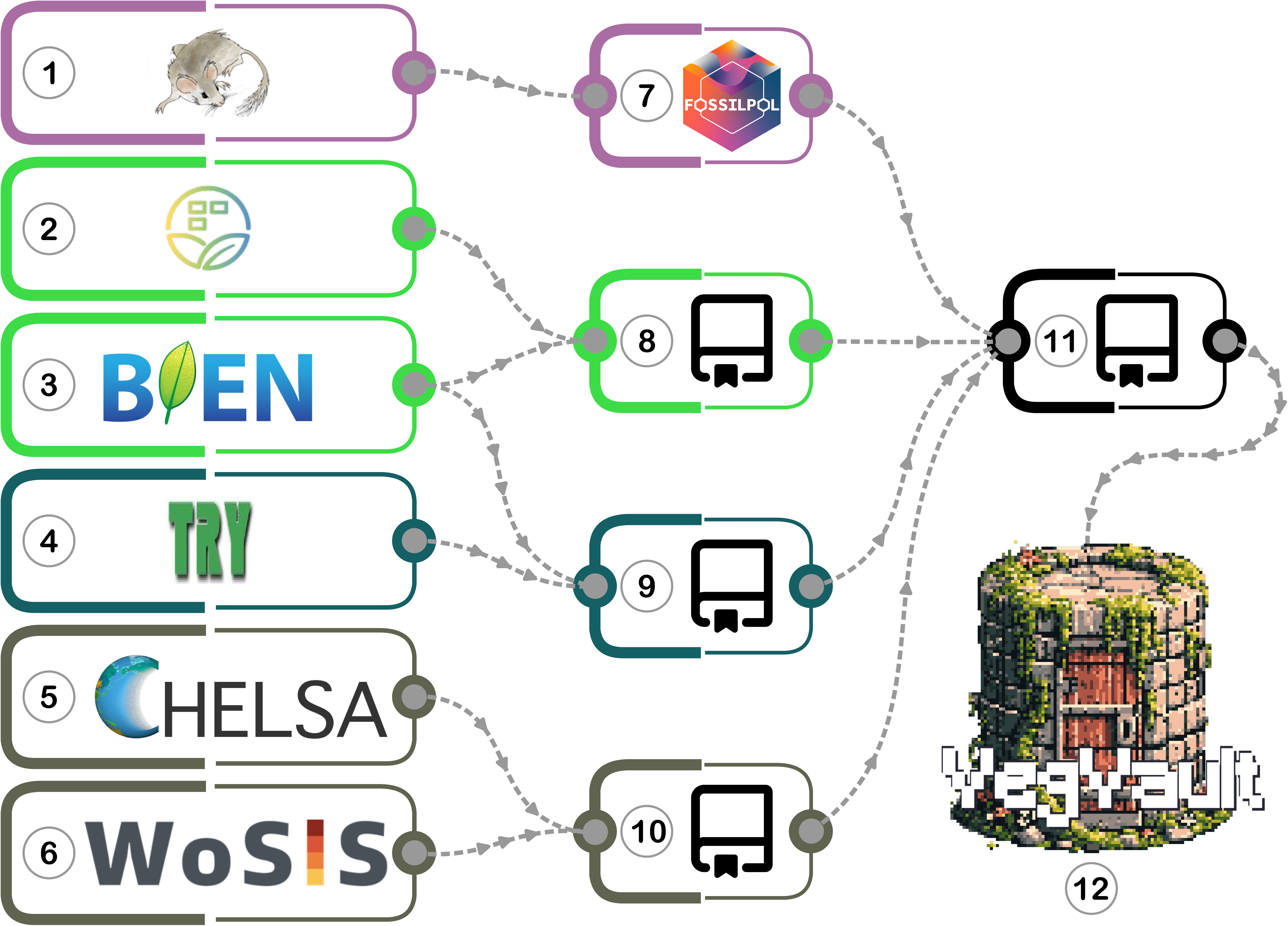
Schematic workflow of accessing and processing all primary data sources into VegVault v1.0.0 database. *1) Neotoma Paleoecological Database*^7^; **2)** *sPlotOpen*^15^; **3)** *Botanical Information and Ecology Network*^6^; **4)** *TRY Plant Trait Database*^8^; 5) *CHELSA*^9^ and *CHELSA-TRACE21*^18,19^; **6)** *World Soil Information Service*^17^; **7)** *FOSSILPOL: Workflow for processing and standardizing global paleoecological pollen data*. A project-specific version with all the settings used for VegVault is stored as GitHub repository *VegVault-FOSSILPOL* (can be accessed as DOI: 10.5281/zenodo.14745060); **8)** The GitHub repository *VegVault-Vegetation_data* used to processes data for contemporary vegetation plots (can be accessed as DOI: 10.5281/zenodo.16729978); **9)** A GitHub repository *VegVault-Trait_data* used to processes data for plant vegetation traits (can be accessed as DOI: 10.5281/zenodo.16729756); **10)** The GitHub repository *VegVault-abiotic_data* used to process climate and soil abiotic data (can be accessed as DOI: 10.5281/zenodo.14745293); **11)** The GitHub repository *VegVault* used to integrate all data together and migrate them into SQLite database (can be accessed as DOI: 10.5281/zenodo.16987345). Additional steps such as the creation of *gridpoints*, grouping of *Traits* into *Trait Domains*, and classification of taxa are also present (for more details see **Methods**); **12)** VegVault - SQLite database (v1.0.0; DOI: 10.48700/datst.t21cz-2nq07); For details about primary data sources, see **Table 1**.

### Data extraction and pre-processing

To ensure the traceability of data, VegVault is designed as a versioned project. Each version (including the current v1.0.0) is created using the git Tags system^32^. Each Tag is a snapshot of code at certain points in time. Each of the individual data pipelines, coded in R programming language, is publicly available and stored in a separate GitHub repository with a specific git Tag and published with DOI (see legend of **Figure 1**).

### Fossil pollen record (Figure 1.7)

Fossil pollen records have been downloaded from the *Neotoma Paleoecological Database*^7^ using their API on 26^th^ June 2023. All data acquisition and processing have been done using the *FOSSILPOL: Workflow for processing and standardizing global paleoecological pollen data*^16^ (version 1.0.0). This includes the selection of depositional environments, ecological groups, chronology control point types, and a minimum number of chronology control points to construct age-depth models. Individual samples and records have been filtered by age limits, number of pollen grains, maximum age interpolation, and number of valid levels. In addition, the accuracy of fossil pollen data is increased by re-estimating all age-depth models using the Bayesian probabilistic approach^33^ and including the information about individual age uncertainty. For field-specific terminology, see Flantua et al.^16^ and for more information about *FOSSILPOL* workflow, see https://bit.ly/FOSSILPOL.

Individual steps in detail are as follows:

- Only records with type “pollen” have been requested with valid geological coordinates. Specifically, longitude between −180 and 180, and latitude between −90 and 90.
- Records have been filtered by their depositional environments, where in this case we only kept lakes, bogs, and mires. See **Table S1** for a list of selected depositional environments.
- Only taxa in specific ecological groups have been accepted. See **Table S2** for a list of selected ecological groups.
- All age-depth models have been recalculated using the {*Bchron*} package^33^. Only records with at least 5 chronology control points of specific types have been retained. See **Table S3** for a list of valid control types.
- To ensure the highest possible data quality and facilitate numerical comparisons across multiple records, several filtering criteria were carefully chosen and applied:

- Exclusion of samples with a total pollen count below a specified threshold. While the preferred minimum number of pollen grains was initially set at 150, this threshold resulted in a significant loss of records, particularly in regions with limited data coverage. To balance data retention and quality, the threshold was adjusted to 125, with the condition that less than 75% of the samples would have a low pollen sum (i.e., lower than 125 pollen grains).
- Exclusion of samples older than 3000 years of the last chronology control point.
- Exclusion of samples beyond 20,000 cal yr BP.
- Exclusion of records if the total number of samples was fewer than 5.

The VegVault-specific version of FOSSILPOL with all the individual R scripts and tables is stored as a GitHub repository titled *VegVault-FOSSILPOL (*https://github.com/OndrejMottl/VegVault-FOSSILPOL/tree/v1.0.0) and can be accessed as DOI: 10.5281/zenodo.14745060.

### Contemporary vegetation plots (**Figure 1**.**8**)

The primary sources of contemporary plot-based vegetation data are *BIEN*^6^ (Botanical Information and Ecology Network) and *sPlotOpen*^15^ (the open-access version of sPlot^15^).

The GitHub repository titled *VegVault-Vegetation_data (*https://github.com/OndrejMottl/VegVault-Vegetation_data/tree/v1.1.0), used to process data for contemporary vegetation plots, can be accessed as DOI: 10.5281/zenodo.16729978.

#### BIEN

The plot-based data have been downloaded using the {*RBIEN*} R package^34^ (version 1.2.7) on 2^nd^ August 2023 by using the function ‘*BIEN::BIEN_plot_datasource()*’. The following columns have been retained: “datasource_id”, “datasource”, “plot_name”, “sampling_protocol”, “methodology_reference”, “methodology_description”, “longitude”, “latitude”, “plot_area_ha”, “subplot”, “individual_count”, “family_matched”, “name_matched”, “name_matched_author”, “verbatim_family”, “verbatim_scientific_name”, “scrubbed_species_binomial”, “scrubbed_taxonomic_status”, scrubbed_family”, “scrubbed_author”. All columns have been renamed using the “snake case”. Finally, rows containing NA values in “datasource_id, “datasource”, “plot_name”, “longitude”, “latitude”, and/or “plot_area_ha” have been filtered out.

#### sPlotOpen

The sPlotOpen dataset^35^ (version 2.0) have been downloaded on 26^th^ September 2023. Tables “DT2.oa” and “header.oa” have been linked via the “PlotObservationID” column, and rows containing NA values in “PlotObservationID”, “GIVD_ID”, “Longitude”, “Latitude”, and/or “Releve_area” have been filtered out. All columns have been renamed using the “snake case”.

### Functional traits of vegetation (**Figure 1.9**)

The primary sources of vegetation functional traits are *TRY Plant Trait Database*^8^ and *BIEN*^6^.

We followed Diaz et al. (2016)^36^ and selected traits representing *Stem specific density*, *Leaf nitrogen content per unit mass*, *Diaspore mass*, *Plant height*, *Leaf area*, and *Leaf mass per area*.

A GitHub repository titled *VegVault-Trait_data* (https://github.com/OndrejMottl/VegVault-Trait_data/tree/v1.2.0), used to process data for plant vegetation traits, can be accessed as DOI: 10.5281/zenodo.16729756.

#### TRY

We made a data request from the *TRY Plant Trait Database* on the 29^th^ of August 2023 (ID of the request is 28498), requesting all values for traits with codes 3106, 4, 3108, 3110, 3112, 3114, 3116, 3117, 14, and 26 (as they are the closest to the Diaz et al. (2016)^36^ description). Next, we used the {*rtry*} R package^37^ (version 1.1.0) to import the data into R.

For each trait (“TraitID”), we extracted all relevant observations (“ObservationID”), ensuring that only observations unique to each trait are considered (i.e., filtering out overlapping observations with other traits). Next, we identified all unique data (“DataID”) associated with each trait. Certain data were excluded (“DataID”: 2221, 2222, 2223, 2224, 2225, 3646, 3647, 3698, 3699, 3727, 3728, 3730, 3731, 3849, 3850, 4029, and 4030) as not meaningful variation of the trait, e.g. *height at 15 days.* We filtered the observations and extracted the actual trait values, ensuring that only non-missing values were kept. Third, we extracted covariate information (additional data associated with each observation but not directly a trait value stored in the “DataName” column). Finally, all columns were renamed using the “snake case”.

As there are differences in trait names across sources of data (e.g., “*Leaf nitrogen (N) content per leaf dry mass*” and “*leaf nitrogen content per leaf dry mass*”), we added a new variable *Trait Domain* that groups traits together following the trait selection by Diaz et al. (2016)^36^. For example, the traits “*Leaf area (in case of compound leaves: leaf, petiole excluded)*” and “*Leaf area (in case of compound leaves: leaf, petiole included)*” are grouped under the “Leaf area” *Trait Domain*. See **Table 3** for the full list of traits and their *Trait Domain*. Such grouping is also aimed to serve as an efficient extraction of traits across both *TRY* and *BIEN*.

**Table 3:**
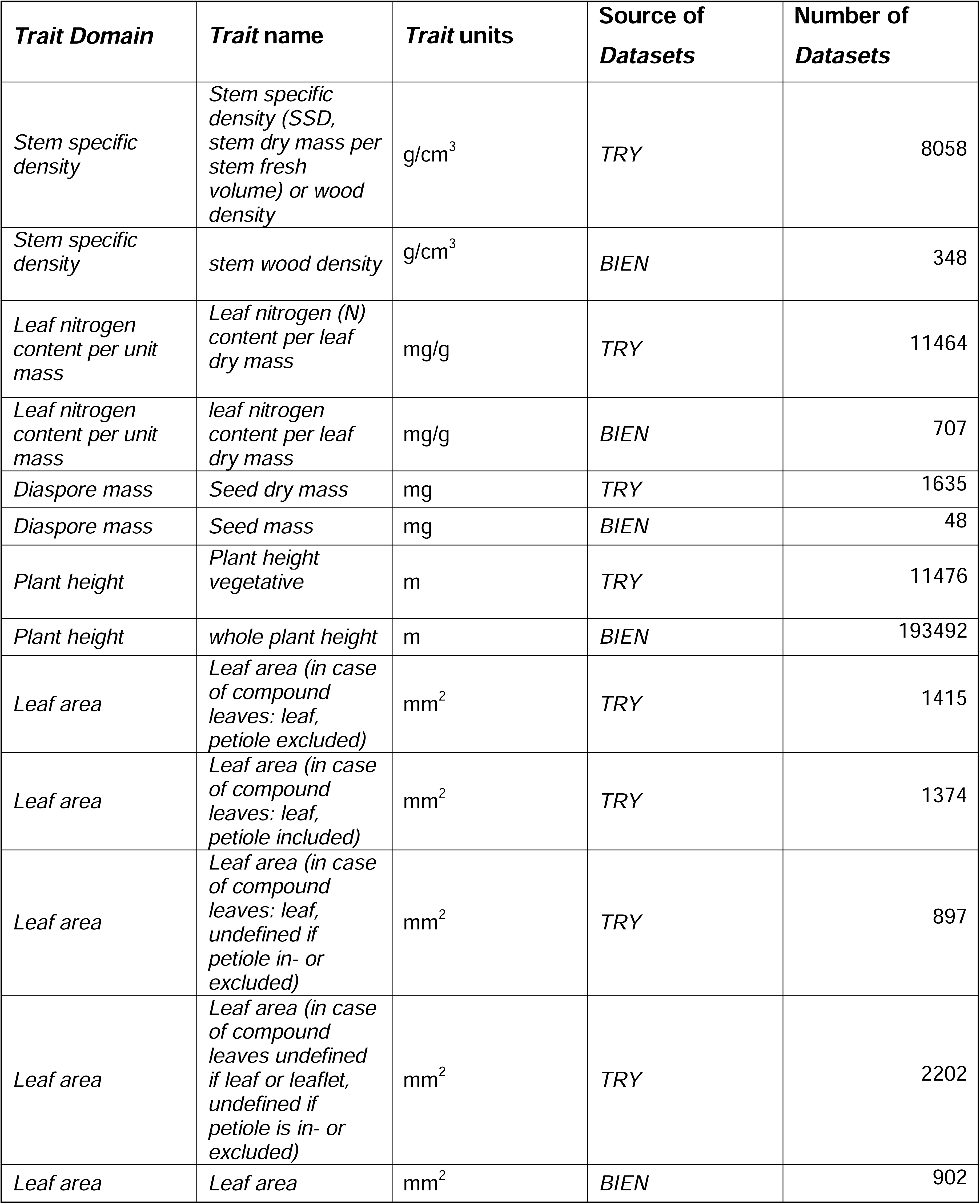
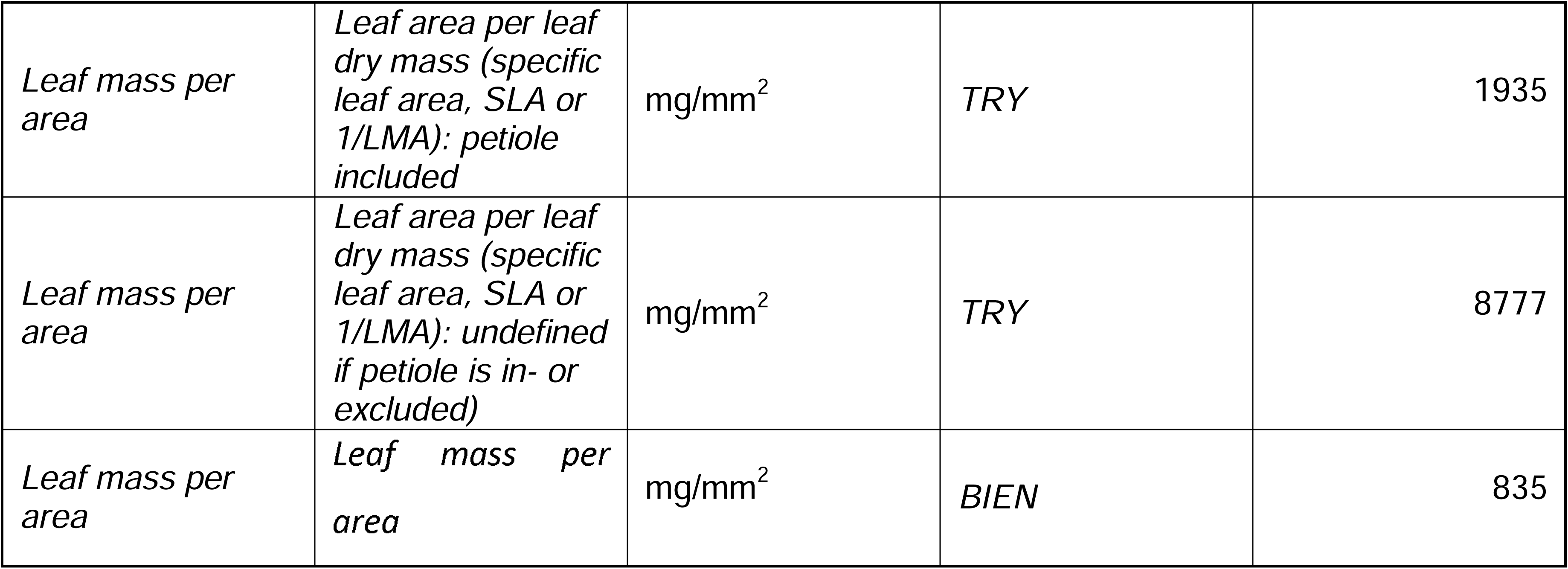
Selected *Traits* presented in the VegVault v1.0.0. The *Traits* were selected based on similarity to Diaz et al. (2016)^36^ and grouped into six main *Trait Domains* to simplify the navigation between different data sources.

#### BIEN

The trait data have been downloaded using the {*RBIEN*} R package^34^ (version 1.2.7) on 15^th^ December 2023 by using the function ‘*BIEN::BIEN_trait_trait()*’. Specifically, we requested data for traits “*whole plant height*”, “stem wood density”, “*leaf area*”, “*leaf area per leaf dry mass*”, “*leaf nitrogen content per leaf dry mass*”, and “*seed mass*”. The following columns have been retained: “trait_name”, “trait_value”, “unit”, “id”, “longitude”, “latitude”, “method”, “url_source”, “source_citation”, “project_pi”, “scrubbed_species_binomial”, and “access”.

### Abiotic data (**Figure 1.10)**

The primary sources of abiotic data are *CHELSA*^9^, *CHELSA-TRACE21*^18,19^, and *WoSIS Soil Profile Database*^17^. The first two data sources provide high-resolution downscaled climatic data, while the latter offers detailed soil information (only available for contemporary data). The abiotic variables included in VegVault are summarised in **Table 4**.

**Table 4:** Overview of abiotic variables present in VegVault v1.0.0.

| Abiotic variable name | Abiotic variable description | Abiotic variable unit | Time period | Source of data | Original spatial resolution | Original temporal resolution |
| --- | --- | --- | --- | --- | --- | --- |
| bio1 | Annual Mean Temperature | °C | contemporary and paleo data | CHELSA & CHELSA-TRACE21 | 1 km (30 arc-second) | 100 years for paleo records |
| bio4 | Temperature Seasonality | °C | contemporary and paleo data | CHELSA & CHELSA-TRACE21 | 1 km (30 arc-second) | 100 years for paleo records |
| bio6 | Min Temperature of Coldest Month | °C | contemporary and paleo data | CHELSA & CHELSA-TRACE21 | 1 km (30 arc-second) | 100 years for paleo records |
| bio12 | Annual Precipitation | kg m-2 year-1 | contemporary and paleo data | CHELSA & CHELSA-TRACE21 | 1 km (30 arc-second) | 100 years for paleo records |
| bio15 | Precipitation Seasonality | Unitless | contemporary and paleo data | CHELSA & CHELSA-TRACE21 | 1 km (30 arc-second) | 100 years for paleo records |
| bio18 | Precipitation of Warmest Quarter | kg m-2 quarter-1 | contemporary and paleo data | CHELSA & CHELSA-TRACE21 | 1 km (30 arc-second) | 100 years for paleo records |
| bio19 | Precipitation of Coldest Quarter | kg m-2 quarter-1 | contemporary and paleo data | CHELSA & CHELSA-TRACE21 | 1 km (30 arc-second) | 100 years for paleo records |
| HWSD2 | Soil type | Unitless | contemporary data | WoSIS | 1 km (30 arc-second) | N/A |

The GitHub repository titled *VegVault-abiotic_data (*https://github.com/OndrejMottl/VegVault-abiotic_data/tree/v1.1.0), used to process climate and soil abiotic data, can be accessed as DOI: 10.5281/zenodo.14745293.

#### CHELSA

Climate data have been downloaded from *CHELSA*^9^ (version 2.1) using the {*ClimDatDownloadR}* R package^38^ on the 8^th^ of September 2023 with the function ‘*ClimDatDownloadR::Chelsa.Clim.download()*’. The specific bio-variables selected are 1, 4, 6, 12, 15, 18, and 19. Next, we used the {*terra*} R package^39^ to spatially aggregate the values using the function ‘*terra::aggregate(factor = 25, fun = “median”)*’.

#### CHELSA-TRACE21

Paleoclimate data have been downloaded from *CHELSA-TRACE21*^18,19^ (version 1.0) on the 31^st^ December 2023. We only downloaded values for each 500-year time-slice between 0 and 18,000 years before present (BP) for the selected bio-variables (1, 4, 6, 12, 15, 18, and 19). For each time slice, we also used the function ‘*terra::aggregate(factor = 25, fun = “median”)*’ to spatially aggregate the values.

#### WoSIS Soil Profile Database

As soil information has been previously confirmed as important predictors for vegetation distribution^40–42^, we opted for including such data, though only representing the present day. We downloaded the soil data from the *WoSIS Soil Profile Database*^17^ on the 11^th^ of September 2023 (both HWSD2_RASTER.zip and HWSD2_DB.zip). We extracted the soil type names (column “HWSD2_SMU_ID”) by combining tables “HWSD2_SMU” and “D_WRB4”, and added this information to the raster. Finally, we resampled the raster using the ‘*terra::resample(method = “near”)*’ function to match the resolution of climate data.

### Data integration (**Figure 1.11**)

All processing pipelines with their corresponding Tags are then migrated into an SQLite database using the GitHub repository titled *VegVault* (https://github.com/OndrejMottl/VegVault/tree/v1.0.0-website_rewiev), which can be accessed as DOI: 10.5281/zenodo.16987345).

### Migrating sPlotOpen vegetation data

The Dataset name (“dataset_name”) has been created from “plot_observation_id” as “splot_[plot_observation_id]”. The original data source (e.g., sub-databases) from column “givd_id” has been stored in the *DatasetSourcesID* table.

The Sample name (“sample_name”) has been created using column “plot_observation_id” as “splot_[plot_observation_id]”. The Sample Size (“sample_size”) has been created from the “releve_area” column. All samples are automatically assigned an age of 0. Taxonomic names have been extracted from column “Species” and the abundances from “Original_abundance”.

### Migrating BIEN vegetation data

The Dataset name (“dataset_name”) has been created as “bien_[row number]”. The original data source (e.g., sub-databases) from column “datasource” has been stored in the *DatasetSourcesID* table. The Sampling method has been extracted from the “methodology_description” column, and its reference has been extracted from the “methodology_description” column.

The Sample name (“sample_name”) has been created as “bien_[row number]”. The Sample Size (“sample_size”) has been created from the “plot_area_ha” column, and multiplied by 10,000 to be stored in square meters. All samples are automatically assigned an age of 0. Taxonomic names have been extracted from column “name_matched” and the abundances from “taxon_name”.

### Migrating fossil pollen data

The Dataset name (“dataset_name”) has been created from “dataset_id” (fossilpol_[dataset_id]). Note that the column “dataset_id” is from the primary source and does not match the “dataset_id” in VegVault. The original data source from column “source_of_data” has been stored in the *DatasetSourcesID* table. The Sampling method has been extracted from the “depositionalenvironment” column. The individual Dataset Reference have been extracted from the column “doi”.

The Sample name (“sample_name”) has been created using columns “dataset_id” and “sample_id” as “fossilpol_[dataset_id]_[sample_id]”. Ages have been extracted from column “levels”.

Age uncertainties from age-depth models have been extracted from “age_uncertainty” column.

Taxonomic names and their abundances have been extracted from the column “counts_harmonised”.

### Migrating TRY functional traits

The Dataset name (“dataset_name”) has been created as “try_[row number]”. The original data source (e.g., sub-databases) from column “dataset” has been stored in the *DatasetSourcesID* table. The individual Dataset Reference have been extracted from the column “reference_source”.

The Sample name (“sample_name”) has been created as “try_[row number]”. The individual Sample reference has been extracted from “dataset_reference_citation”. All samples are automatically assigned an age of 0.

Trait names have been extracted from “trait_full_name”, taxonomic names from the column “acc_species_name”, and trait values from “trait_value”.

### Migrating BIEN functional traits

The Dataset name (“dataset_name”) has been created as “bien_traits_[row number]”. The original data source (e.g., sub-databases) from column “project_pi” has been stored in the *DatasetSourcesID* table. The individual Dataset Reference have been extracted from the column “source_citation”.

The Sample name (“sample_name”) has been created using column “id” as “bien_traits_[id]”. All samples are automatically assigned an age of 0.

Trait names have been extracted from “trait_name”, taxonomic names from the column “scrubbed_species_binomial”, and trait values from “trait_value”. Trait “leaf mass per area” is calculated from “leaf area per leaf dry mass” as 1/value.

### Creation of *gridpoints* of abiotic data

We developed a data structure that provides readily available environmental context for each vegetation (and trait) record by creating spatio-temporal links between these records and abiotic information. As the rasters are not suitable to be stored in an SQLite database, we created artificial points, called ‘*gridpoints’*, located in the middle of each raster cell. This resulted in a uniform spatio-temporal matrix of gridpoints holding the abiotic information (see **Figure 2B** for an example area).

**Figure 2.**
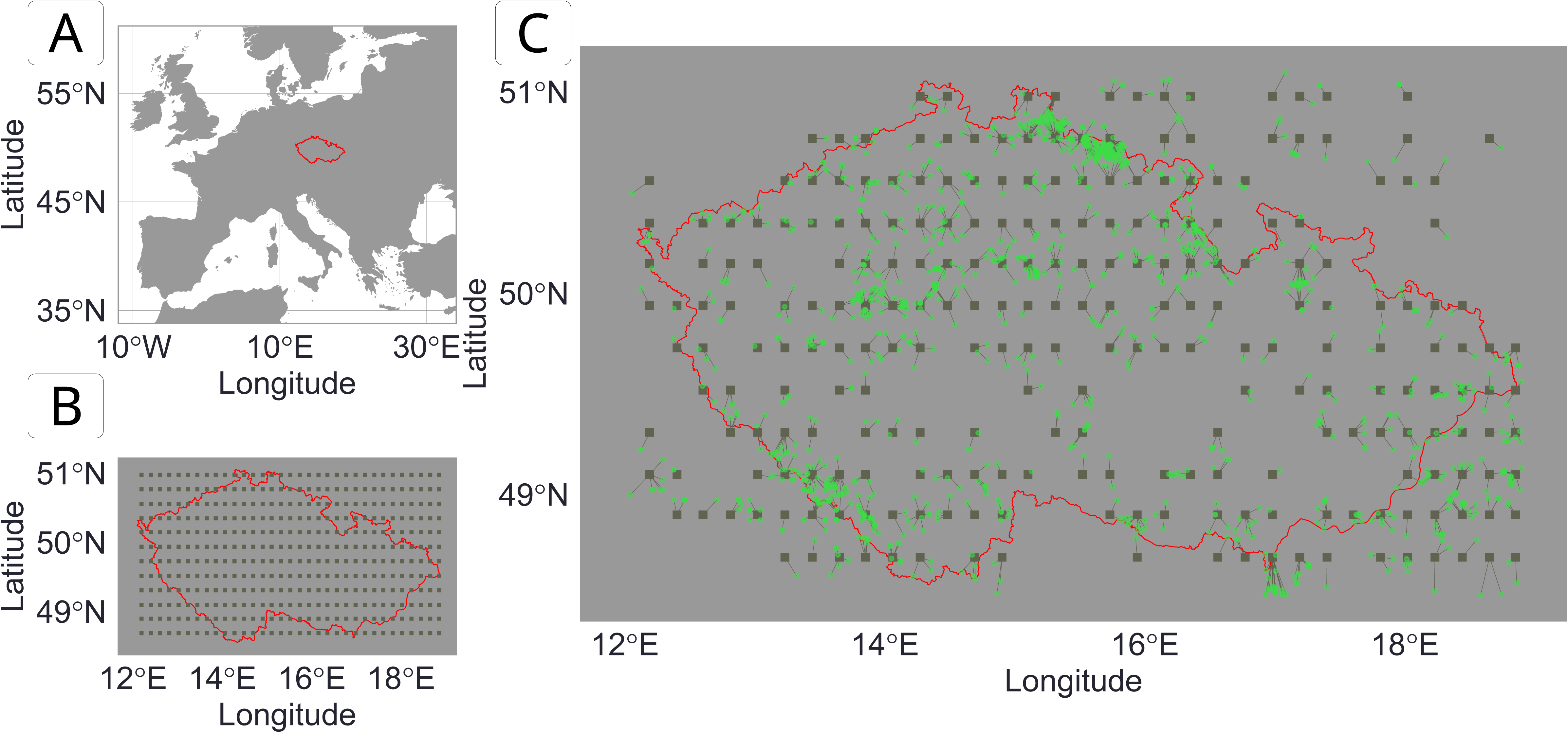
Example of the resolution of the *gridpoint Datasets* extracting abiotic information. A: Map of Europe. The red line indicates country borders with the Czech Republic as the focus of the map. B: Distribution of the individual *gridpoints* (grey squares) within the Czech Republic (red). C: Linking vegetation plots and *gridpoints*. The green circles represent the contemporary vegetation plots. The grey lines connect the vegetation points to their closest *gridpoint*. Note that the abiotic data is provided to each vegetation plot from their closest *gridpoint* (i.e., the vegetation plots connected to the same gridpoint will have the same abiotic data). *Gridpoints* without vegetation plots in their spatio-temporal vicinity (here contemporary data) were excluded from the VegVault to reduce the database size. See Methods section for more details.

Gridpoints Dataset names (“dataset_name”) have been created as “geo_[longitude]_[latitude]” and gridpoints Sample names (“sample_name”) have been created as “geo_[longitude]_[latitude]_[age]”. Next, we linked *gridpoints* and other non-*gridpoint Samples*, namely *vegetation_plot*, *fossil_pollen_archive*, and *traits* (see **Internal Dataset Structure** section), and calculated the spatial and temporal distances between them. We kept any *gridpoint Sample* within 50 km and/or 5000 years from any other non-*gridpoint Sample* and discarded the rest. See **Figure 2C** for an example of linking abiotic information at *gridpoints* and vegetation data. In VegVault, users can select the information for each non-*gridpoint Sample* from the closest spatio-temporal abiotic data or get the average from all surrounding *gridpoints*. We hope that by providing such a comprehensive and well-linked structure between vegetation and abiotic data, VegVault enhances the ability to study the interactions between vegetation and their environment, facilitating the workflow for advanced ecological research and modelling efforts.

### Taxon classification

As VegVault consists of data on taxa from various sources, the {*taxospace*} R package^21^ has been used to classify the diverse taxa into a unifying backbone in the VegVault. The {*taxospace*} tool automatically aligns taxon names with the taxonomical backbone of the *GBIF*^20^. Specifically, we find the best match of the raw names of taxa using the *Global Names Resolver*^43^, which is then aligned with *GBIF*. The resulting information of taxonomic classification, detailed up to the family level, is stored for each taxon, ensuring consistency and facilitating comparative analyses across different datasets. However, note that taxonomic classification down to the species level is not available for each taxon (e.g., some fossil pollen types can only be identified to the genus or family level). For several taxa, we were unable to find a matching classification. Note that the taxon classification is additional information; the original taxon name is always present and returned by default. Finally, we would like to stress to the user that this classification is an automated process and may contain errors.

### Data Records

The VegVault v1.0.0 is available at the *National Repository*^44^ (‘*REPO*’) under DOI <u>10.48700/datst.t21cz-2nq07</u>. It is an SQLite database of a size of ∼ 110 GB, which contains > 480,000 *Datasets*, > 13,000,000 *Samples*, > 100,000 *Taxa*, 6 vegetation *Traits*, > 11,000,000 *Trait* values, and 8 abiotic variables. The spatio-temporal overview of all vegetation data can be found in **Figure 3**.

**Figure 3.**
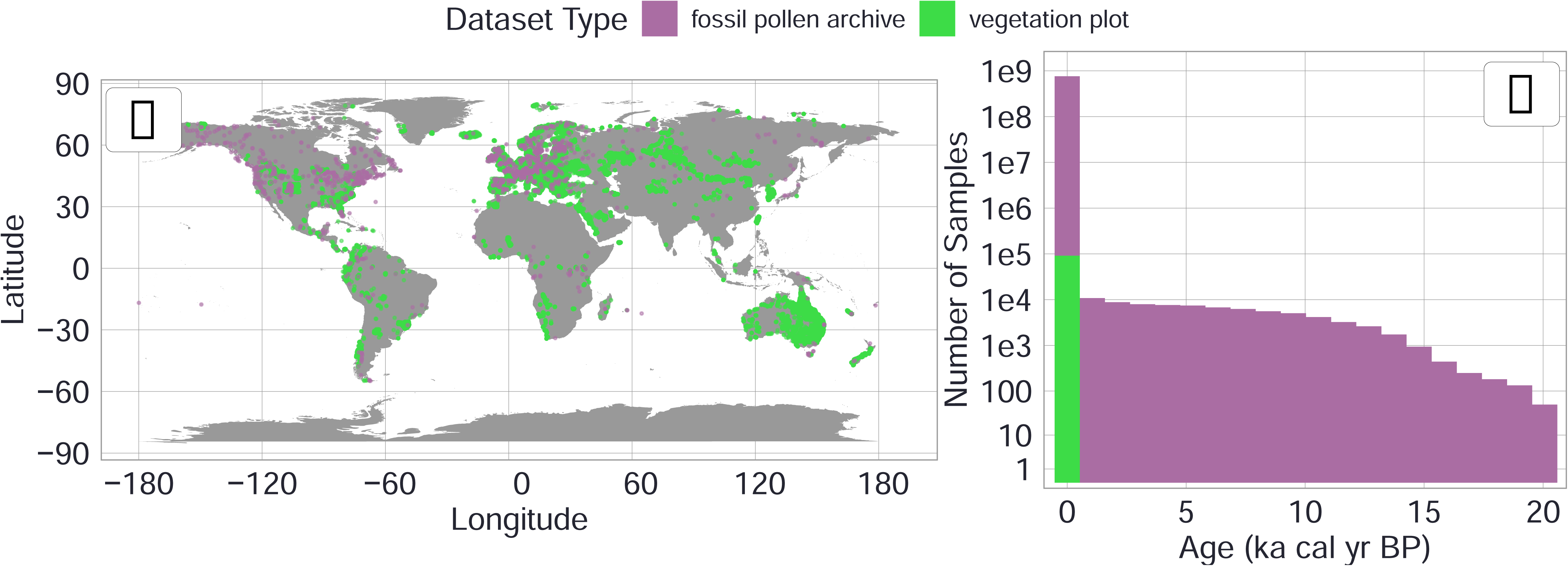
Spatio-temporal distribution of vegetation Samples (for both vegetation plots and fossil pollen archives). Note that this is an overview of the current state of VegVault (v1.0.0.). A: Spatial distribution with each point representing one plot. B: Temporal distribution of individual Samples.

### Internal Dataset Structure

Currently, the dataset consists of 31 interconnected tables with 87 fields (variables), which are described in detail below. See **Figure 4** for the structure (schema) of the SQLite database. However, the internal structure of VegVault may not be of relevance to a user, as the {*vaultkeepr*} R package processes all the data to output only the most relevant information in a “ready-to-analyse” format (see **Usage Notes** section). We will describe all tables and their inter-relationships in the following sections.

**Figure 4:**
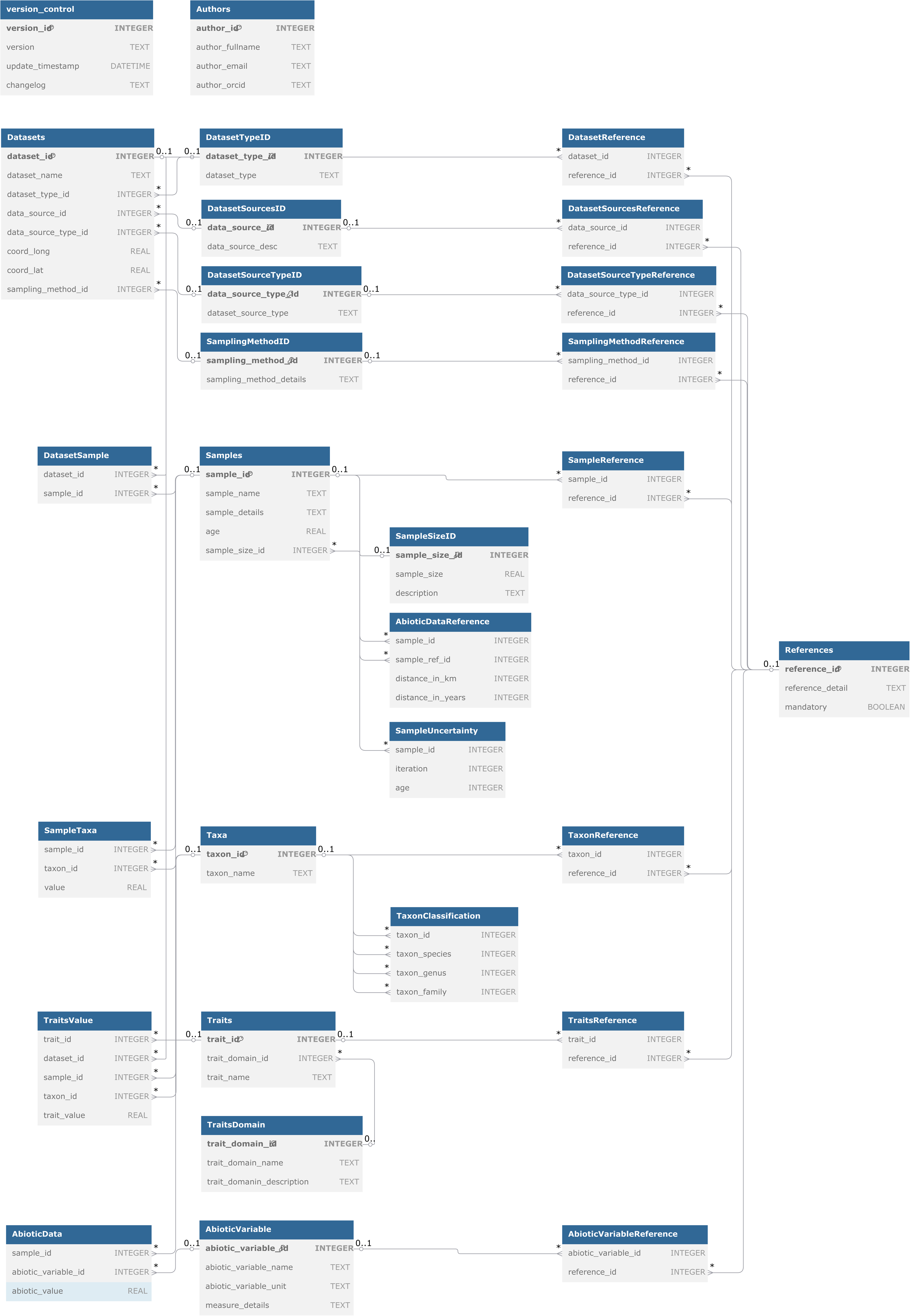
Visualisation of database structure (schema) created using https://dbdiagram.io/. Each table contains its name, the names of individual columns, and the data type of each column. In addition, primary key columns are indicated using a small key icon. Individual lines indicate the connections

### Metadata tables

There are currently several tables that are not directly linked to the data. First is the *Authors* table, which describes the authors and maintainers of VegVault, including the contact information. The next table is *version_control* table, which stores the information about database versions with a description of individual changes at a specific point in time. See **Table 5** and **Table 6** for the description of individual columns, respectively. Finally, there are *sqlite_stat1* and *sqlite_stat4* tables, which hold descriptive information about database indexes, used for improving the performance of data extractions. See **Table 7** and **Table 8** for the description of individual columns, respectively.

**Table 5:** Column names and types for table *Authors*.

| Column name | Data type | Description |
| --- | --- | --- |
| author_id | integer | ID of an Author (unique) |
| author_fullname | character | Full name of an *Author* |
| author_email | character | Contact email of an *Author* |
| author_orcid | character | ORCID ID of an *Author* |

**Table 6:** Column names and types for table *version_control*.

| Column name | Data type | Description |
| --- | --- | --- |
| id | integer | ID a database version (unique) |
| version | character | Version number |
| update_date | character | Date of the creation of that version |
| changelog | character | Text description of main changes in the database |

**Table 7:** Column names and types for table *sqlite_stat1*.

| Column name | Data type | Description |
| --- | --- | --- |
| tbl | character | Name of a table |
| idx | character | Name of a database index |
| stat | character | Statistical information. Specifically, comma-separated list of numbers. |

**Table 8:** Column names and types for table *sqlite_stat4*.

| Column name | Data type | Description |
| --- | --- | --- |
| tbl | character | Name of a table |
| idx | character | Name of a database index |
| neq | character | Number of entries in the index equal to the sample key |
| nlt | character | Number of entries less than the sample key |
| ndlt | character | Number of distinct values less than the sample key |
| sample | character | A sample value from the index (binary-encoded key values) |

### *Datasets* table

The table contains one *Dataset* per row, with each *Dataset* may contain: a unique *Dataset ID* (“dataset_id*”*), *Dataset name* (“dataset_name”), geographic location (“coord_lat”, “coord_long”), Dataset Type (“dataset_type_id*”), Dataset Source* (“data_source_id”), *Dataset Source Type* (“dataset_source_type_id”), and *Sampling Method* (“sampling_method_id”). See **Table 9** for the description of individual columns.

**Table 9:** Column names and types for table *Datasets*.

| Column name | Data type | Description |
| --- | --- | --- |
| dataset_id | integer | ID of a *Dataset* (unique) |
| dataset_name | character | Name of each *Dataset* |
| dataset_type_id | integer | ID of a *Dataset Type* |
| data_source_id | integer | ID of a Dataset Source |
| data_source_type_id | integer | ID of a Dataset Source-Type |
| coord_long | numeric | Geographical coordinates – longitude |
| coord_lat | numeric | Geographical coordinates – latitude |
| sampling_method_id | integer | ID of a *Sampling Method* |

### DatasetTypeID table

The table defines the most basic classification of each *Dataset*, ensuring that the vast amount of data is categorised systematically (see **Table 10** for the description of individual columns). Currently, VegVault contains the following *Dataset Types*:

- ***’vegetation_plot’***: This type includes contemporary vegetation plot data, capturing contemporary vegetation characteristics and distributions.
- ***’fossil_pollen_archivè***: This type encompasses past vegetation plot data derived from fossil pollen records, providing insights into past vegetation and climate dynamics.
- ***’traits’***: This type contains functional trait data, detailing specific characteristics of plant species that influence their ecological roles.
- ***’gridpoints’***: This type holds artificially created *Datasets* to manage abiotic data, here climate and soil information (a dataset type created to hold abiotic data, see details in the **Methods** section).

**Table 10:** Column names and types for table *DatasetTypeID*.

| Column name | Data type | Description |
| --- | --- | --- |
| dataset_type_id | integer | ID of a *Dataset Type* (unique) |
| dataset_type | character | Text description of individual *Dataset Types* (currently *vegetation_plot*, *fossil_pollen_archive*, *traits*, *gridpoints*) |

See **Figure 5A** for an overview of the number of *Datasets* in each *Dataset Type*.

**Figure 5.**
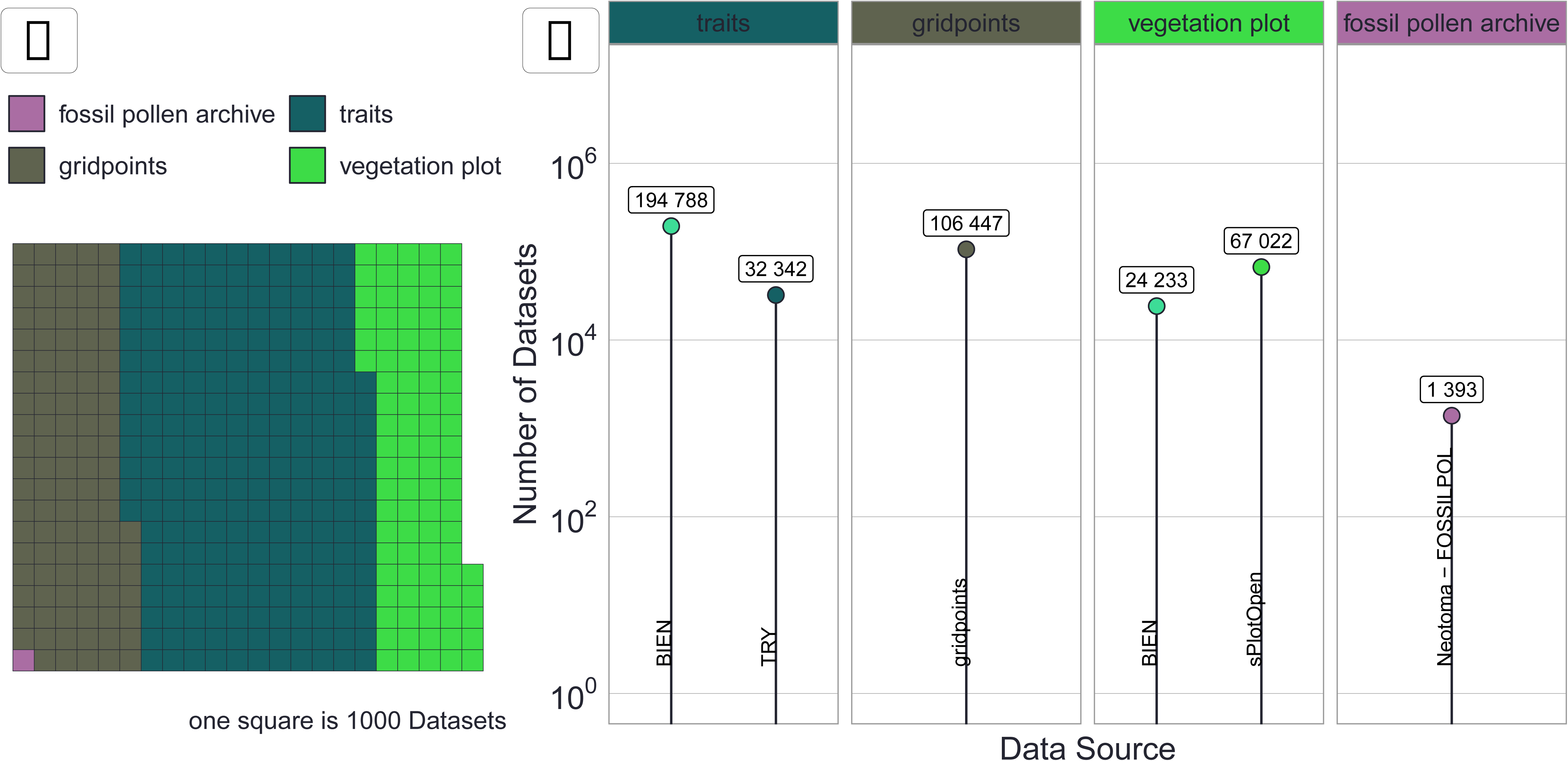
Visualisation of the number of *Datasets*. A: Waffle plot coloured by *Dataset Type* with each square representing ∼1000 *Datasets*. B: Number of *Datasets* by *Dataset Source-Type* grouped by *Dataset Type*. See Internal Dataset Structure section for more details.

### DatasetSourceTypeID table

The table contains information about the primary data source (see **Table 11** for the description of individual columns). Therefore, each *Dataset* is derived from a specific *Source-Type* including:

- ***BIEN***^6^ (Botanical Information and Ecology Network).
- ***sPlotOpen***^15^ (The open-access version of sPlot).
- ***TRY***^8^ (TRY Plant Trait Database).
- ***Neotoma-FOSSILPOL*** (FOSSILPOL^16^: Workflow for processing and standardizing global paleoecological pollen data from *Neotoma*^7^) Note that we specifically state *Neotoma*-*FOSSILPOL* and not just *Neotoma*, as *FOSSILPOL* not only provides the data acquisition but also alters it (e.g., creating new age-depth models). It also addresses major challenges in paleoecological data integration, such as age uncertainty, by incorporating probabilistic age-depth models and their associated uncertainty matrices. This enables the propagation of temporal uncertainty in subsequent analyses, a critical advancement for robust macroecological studies, previously flagged as a major issue with paleo-data^45^.
- ***gridpoints*** (artificially created *Datasets* for abiotic data). See details in the **Methods** section.

**Table 11:** Column names and types for table *DatasetSourceTypeID*.

| Column name | Data type | Description |
| --- | --- | --- |
| data_source_type_id | integer | ID of a *Dataset Source-Type* (unique) |
| dataset_source_type | character | Text description of individual *Dataset Source-Type* (currently, *BIEN*, *sPlotOpen*, *TRY*, *Neotoma-FOSSILPOL*, *gridpoints*) |

See **Figure 5B** for an overview of the number of *Datasets* for each *Dataset Source-Type*.

### DatasetSourceTypeReference table

The table links the *Dataset Source-Types* to their specific *Reference*. Note that one *Dataset Source-Type* may have several *References. See* **Table 12** for the description of individual columns.

**Table 12:** Column names and types for table *DatasetSourceTypeReference*.

| Column name | Data type | Description |
| --- | --- | --- |
| data_source_type_id | integer | ID of a *Dataset Source-Type* |
| reference_id | integer | ID of a *Reference* |

### DatasetSourcesID *table*

The table contains information about the provider (*Dataset Source*) of the original data for the primary data source. The VegVault v1.0.0 currently includes 706 sources of *Datasets,* where each dataset can also have one or more direct references to specific data (see below), ensuring that users can accurately cite and verify the sources of their data. See **Table 13** for the description of individual columns.

**Table 13:** Column names and types for table *DatasetSourcesID*.

| Column name | Data type | Description |
| --- | --- | --- |
| data_source_id | integer | ID of a *Dataset Source* (unique) |
| data_source_desc | character | Text description of individual *Dataset Sources* (e.g., name of the sub-database from the primary source) |

### DatasetSourcesReference table

The table links the *Dataset Sources* to their specific *Reference*. Note that one *Dataset Source* may have several *References.* See **Table 14** for the description of individual columns.

**Table 14:** Column names and types for table *DatasetSourcesReference*.

| Column name | Data type | Description |
| --- | --- | --- |
| data_source_id | integer | ID of a *Dataset Source* |
| reference_id | integer | ID of a *Reference* |

### SamplingMethodID table

The table contains information about specific *Sampling Method* in the form of an abbreviated ID, which is only available for both *vegetation_plot* and *fossil_pollen_archive Datasets*, and varies substantially, reflecting the diverse nature of the data collected. Such information is crucial for understanding the context and limitations of each *Dataset Type*. For contemporary vegetation plots, sampling involves standardised plot inventories and surveys that capture detailed vegetation characteristics across various regions. Fossil pollen data are collected from sediment records from numerous different depositional environments^16^ representing past vegetation and climate dynamics. See **Table 15** for the description of individual columns.

**Table 15:** Column names and types for table *SamplingMethodID*.

| Column name | Data type | Description |
| --- | --- | --- |
| sampling_method_id | integer | ID of a *Dataset Sampling Method* (unique) |
| sampling_method_details | character | Text description of individual *Dataset Sampling Method* (e.g., name of the sub-database from the primary source) |

### SamplingMethodReference table

The table links the *Dataset Sampling Methods* to their specific *Reference*. Note that one *Dataset Sampling Methods* may have several *References.* See **Table 16** for the description of individual columns.

**Table 16:** Column names and types for table *SamplingMethodReference*.

| Column name | Data type | Description |
| --- | --- | --- |
| sampling_method_id | integer | ID of a *Dataset Sampling Method* |
| reference_id | integer | ID of a *Reference* |

### DatasetReference table

This table links individual *Datasets* to their specific *References* (see **Table 17** for the description of individual columns). Note that one *Dataset* may have several *References*. To support robust and transparent scientific research, each *Dataset* in VegVault can have multiple *References* at different levels as the raw data was pooled from other databases that provide data from various authors. The *Dataset Source-Type*, *Dataset Source*, and *Sampling Method* can all have their own *References*, providing detailed provenance and citation information. This multi-level referencing system enhances the traceability and validation of the data.

**Table 17:**
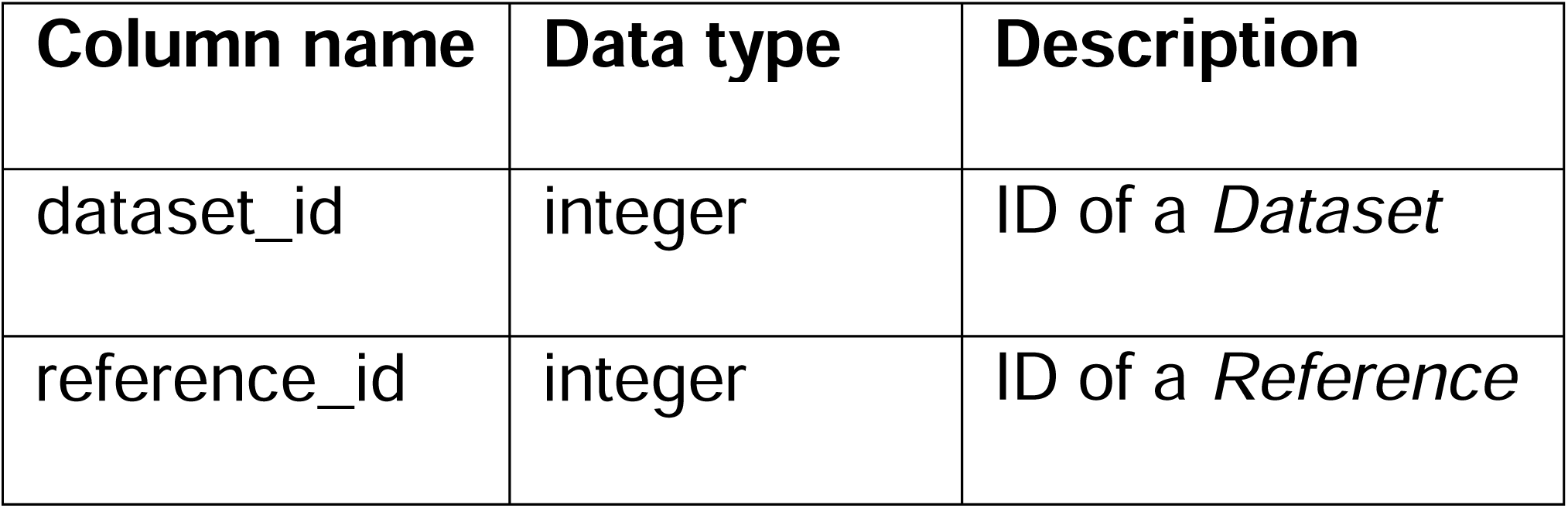
Column names and types for table *DatasetReferences*.

| Column name | Data type | Description |
| --- | --- | --- |
| dataset_id | integer | ID of a *Dataset* |
| reference_id | integer | ID of a *Reference* |

### *Samples* table

*Samples* represent the main unit of data in the *VegVault*, serving as the fundamental building blocks for all analyses. There are currently over 13 million *Samples* in the VegVault v1.0.0 (of which ∼ 1.6 million are *gridpoints* of abiotic data, see **Methods** section). For contemporary and fossil pollen records, each Sample is a representation of vegetation community at a specific location and time. The *Samples* table contains one *Sample* per row, with each *Sample* may contain: a unique *Sample ID* (“sample_id*”*), *Sample name* (“sample_name”), temporal information about *Sample* (“age*”*), *Sample size* (size of the plot if available; “sample_size_id”), and additional information about *Sample* (“sample_details”; this is currently not being used in v1.0.0.). As VegVault encompasses both contemporary and paleo-data, accurate age information for each *Sample* is needed. Contemporary *Samples* are assigned an age of 0, while *Samples* from fossil pollen records are in calibrated years before the present (cal yr BP). The ‘present’ is here specified as 1950 AD. *See* **Table 18** for the description of individual columns.

**Table 18:** Column names and types for table *Samples*.

| Column name | Data type | Description |
| --- | --- | --- |
| sample_id | integer | ID of a *Sample* (unique) |
| sample_name | character | Name of a Sample |
| sample_details | character | Specific description of a *Sample*. Currently not being used. |
| age | numeric | Age of sample. Mainly used for *fossil_pollen_archives*, where note the age of a *Sample* in calibrated years before present. Note that all contemporary *Samples*, have age of 0. |
| sample_size_id | integer | ID of a *Sample Size* |

### DatasetSample table

Each *Sample* is linked to a specific *Dataset* via the *DatasetSample* table, which ensures that every *Sample* is correctly associated with its corresponding *Dataset Type* (whether it is *vegetation_plot*, *fossil_pollen_archive*, *traits*, or *gridpoint*) and other *Dataset* properties (e.g., geographic location). One *Dataset* contains several *Samples* only in cases where *Samples* differ in time (“age*”*). See **Table 19** for the description of individual columns.

**Table 19:**
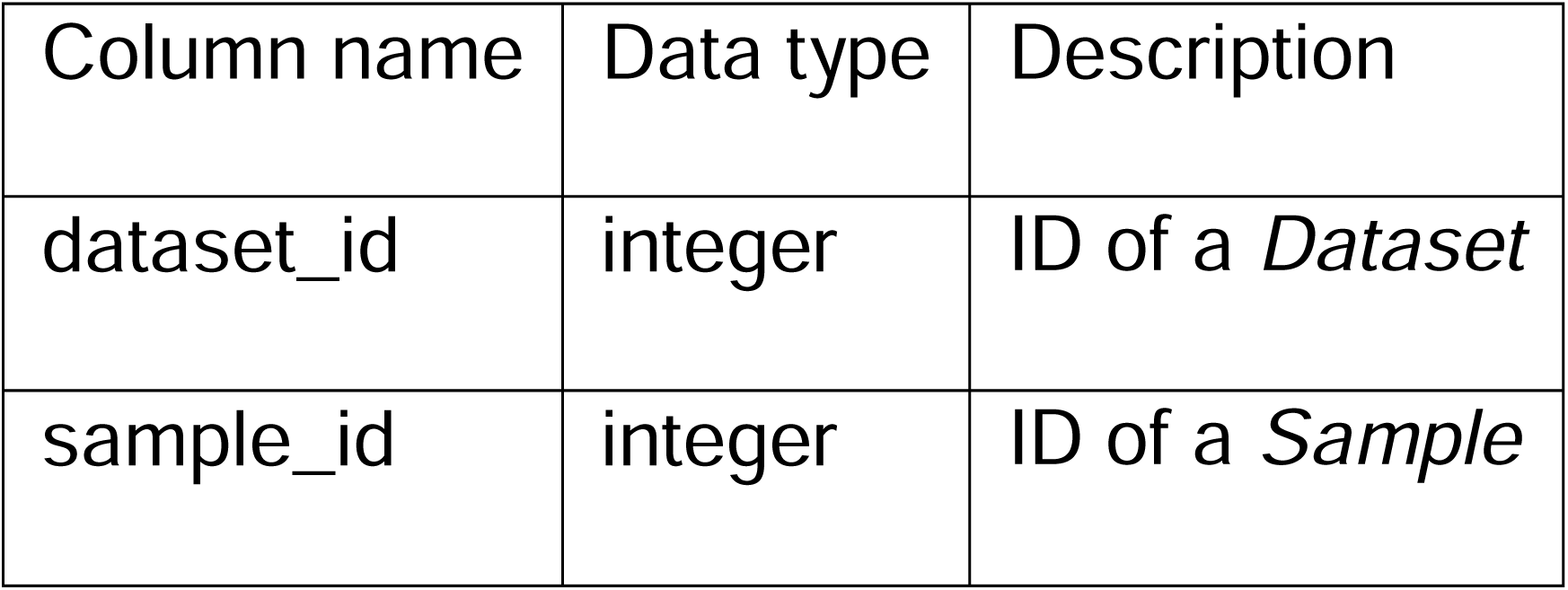
Column names and types for table *DatasetSample*.

| Column name | Data type | Description |
| --- | --- | --- |
| dataset_id | integer | ID of a *Dataset* |
| sample_id | integer | ID of a *Sample* |

### SampleSizeID table

The size of vegetation plots can vary substantially. This detail is crucial for ecological studies where plot size can influence species diversity and abundance metrics, thus impacting follow-up analyses and interpretations^46^. To account for this variability, information about the plot size is recorded separately for each contemporary *Sample*. See **Figure 6** for an overview of the number of *Samples* in each size category and **Table 20** for the description of individual columns.

**Figure 6.**
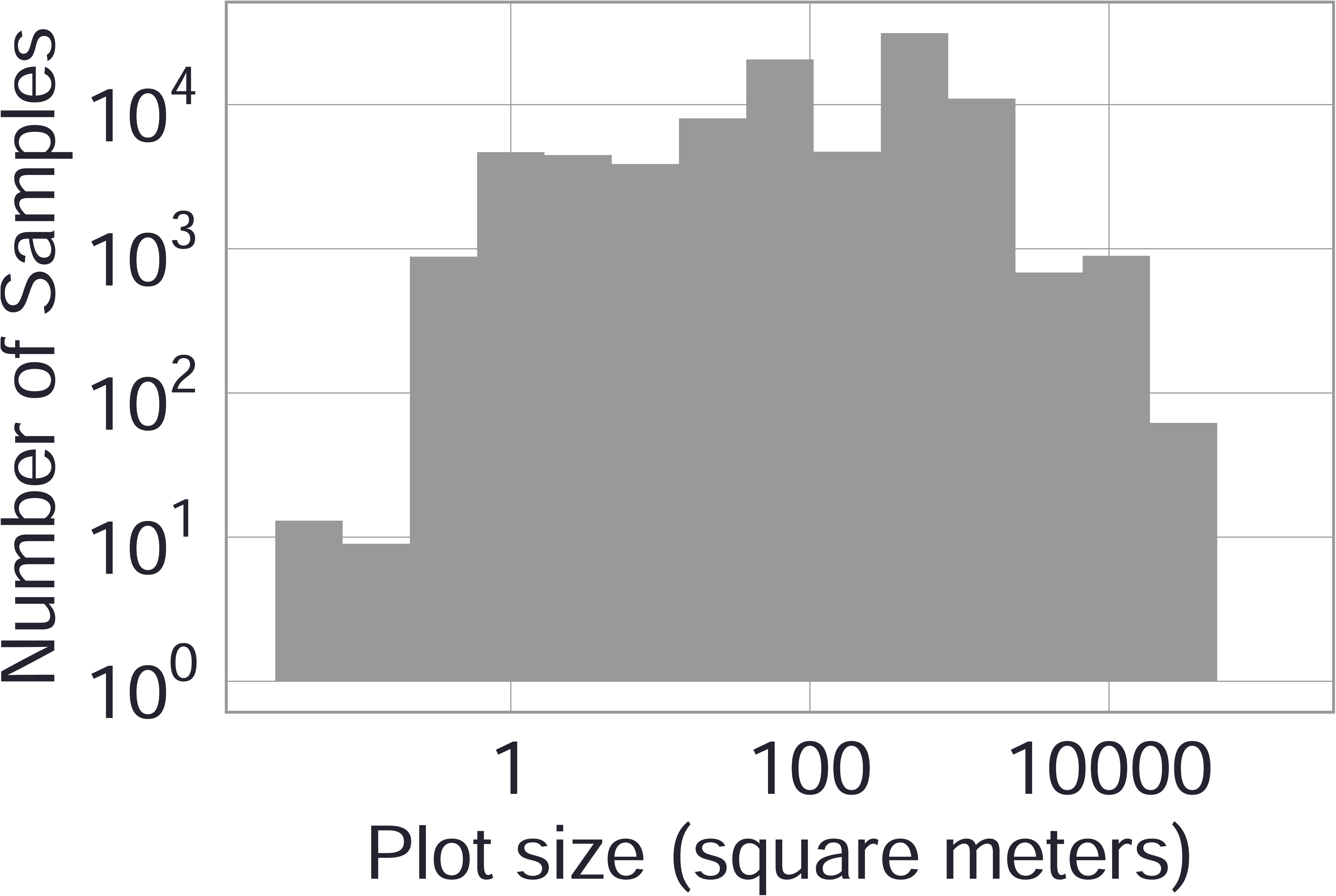
Distribution of *Samples* by vegetation plot sizes. In VegVault, only *vegetation_plot Samples* and *Datasets* have such information. See Internal Dataset Structure section for more details.

**Table 20:** Column names and types for table *SampleSizeID*.

| Column name | Data type | Description |
| --- | --- | --- |
| sample_size_id | integer | ID of a size category (unique) |
| sample_size | numeric | Numeric expression of size |
| description | character | Mostly description of units in which the values are stored |

### SampleUncertainty table

Each *Sample* from the *fossil_pollen_archive Dataset* is also associated with an uncertainty matrix generated during the re-estimation of ages using FOSSILPOL workflow^16^. This matrix provides a range of potential ages derived from age-depth modelling, reflecting the inherent uncertainty in dating paleoecological records. See an example of detailed age uncertainty information for one fossil pollen record in **Figure 7**, and the **Table 21** for the description of individual columns

**Figure 7:**
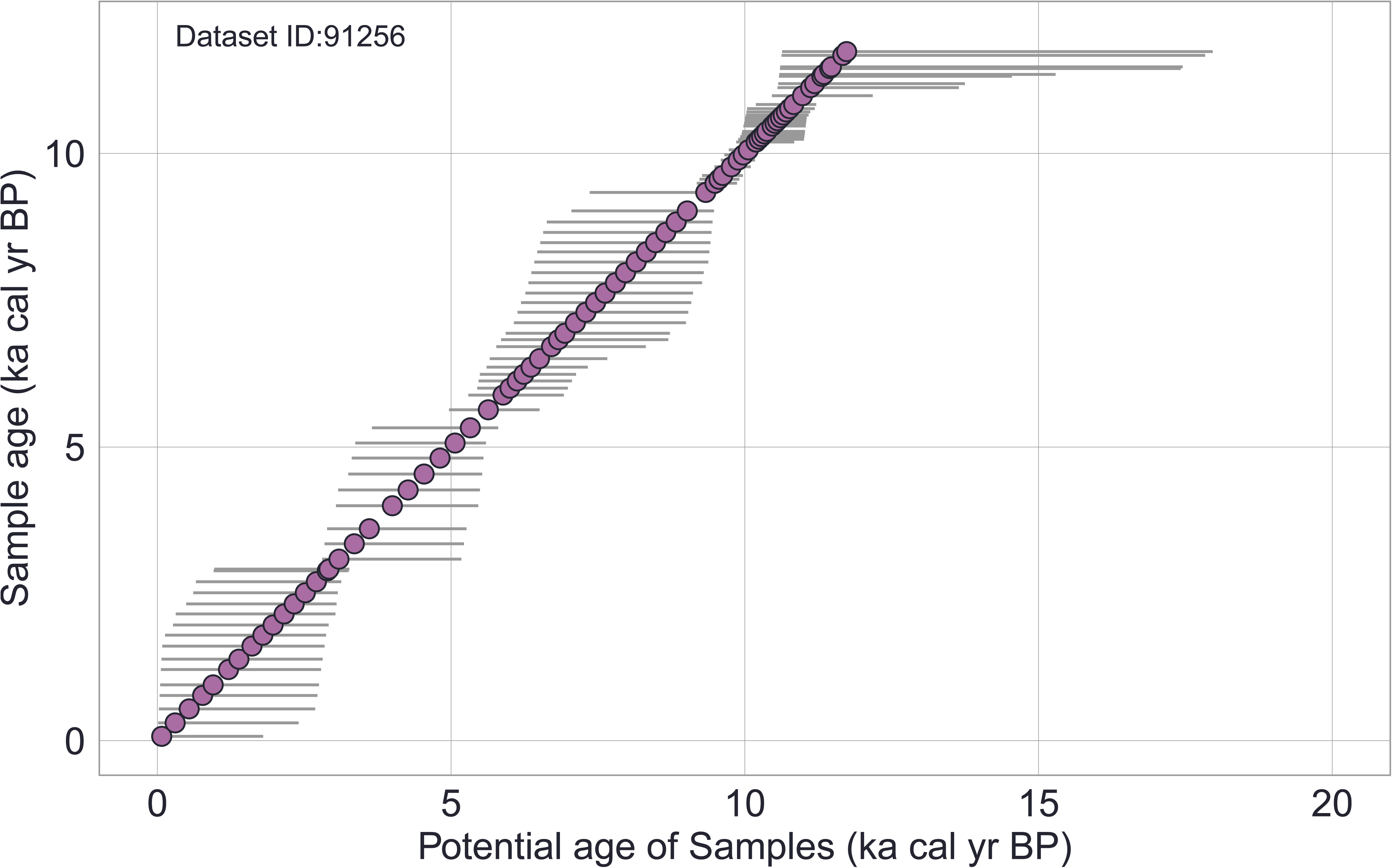
Example of age uncertainty of Samples for selected fossil pollen Dataset (ID: 91256). Each point represents one *Sample*. Each horizontal line represents a range of all potential ages associated with that *Sample*. Potential ages are gathered from age-depth models and stored in the *SampleUncertainty* table. See **Internal Dataset Structure** section for more details about the data structure and *FOSSILPOL* workflow^16^ for more details about age uncertainty from age-depth models.

**Table 21:** Column names and types for table *SampleUncertainty*.

| Column name | Data type | Description |
| --- | --- | --- |
| sample_id | integer | ID of a *Sample* |
| iteration | integer | ID of a iteration from age depth model. Currently, the is 1000 iteration per each Sample. |
| age | integer | Potential age of a Sample |

### SampleReference table

Each *Sample* in VegVault can have specific *References* in addition to those at the *Dataset*-level. These individual *Sample References* provide detailed provenance and citation information, ensuring that users can trace the origin and validation of each data point. Note that a single *Sample* can have several *References*. This level of referencing enhances the transparency and reliability of the data, especially when the dataset continues to be updated in the future. See **Table 22** for the description of individual columns.

**Table 22:** Column names and types for table *SampleReference*.

| Column name | Data type | Description |
| --- | --- | --- |
| sample_id | integer | ID of a *Sample* |
| reference_id | integer | ID of a *Reference* |

**Table 23:** Column names and types for table *Taxa*.

| Column name | Data type | Description |
| --- | --- | --- |
| taxon_id | integer | ID of a *Taxon* (unique) |
| taxon_name | character | Name of a *Taxon* from primary source. |

### *Taxa* table

The VegVault database records the original taxonomic names derived directly from the primary data sources, and currently, it holds over 100 thousand taxonomic names. See **Methods** for the names of columns used from the primary sources. See **Table 19** for the description of individual columns.

### SampleTaxa table

Each individual *Taxon* is linked to corresponding *Samples* through the *SampleTaxa* table, ensuring accurate and systematic association between species and their ecological data. Note that the abundance information varies across the primary data sources (see **Methods** for the names of columns used as abundance). Therefore, users have to be careful while processing data from various sources. See **Table 24** for the description of individual columns.

**Table 24:** Column names and types for table *SampleTaxa*.

| Column name | Data type | Description |
| --- | --- | --- |
| sample_id | integer | ID of a *Sample* |
| taxon_id | integer | ID of a *Taxon* |
| value | numeric | Abundance representation of a *Taxon* (the units may differ among primary sources, i.e., *Dataset Source-Types*) |

### TaxonClassification table

Each taxonomic name undergoes an automated classification (see **Methods** section), and results are stored in the *TaxonClassification* table. Taxonomic classification for some *Taxa* might be only available down to the genus or family level, while some most of the data is classified to species level. See **Figure 8** for an overview of the number of species, genus and family classification. However, 1312 (1.2%) of all *Taxa* currently do not contain classification, and only their name from the primary source is available. The best possible name is returned for each request, which means that *Taxa* without classification will return the name from the primary source. See **Table 25** for the description of individual columns.

**Figure 8.**
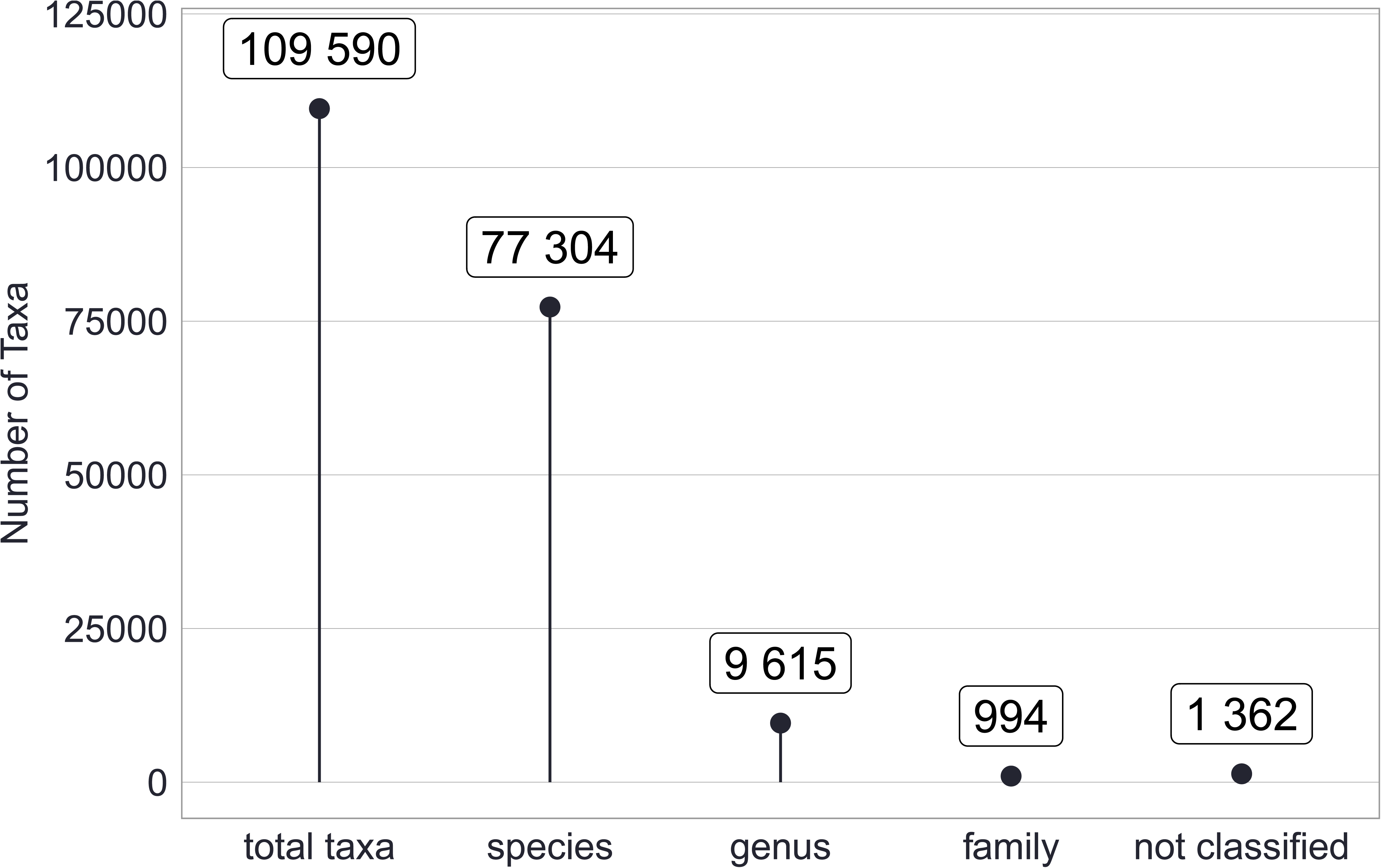
Total number of *Taxa* and their classification present in VegVault. See Methods section for more information about the classification process.

**Table 25:** Column names and types for table *TaxonClassification*.

| Column name | Data type | Description |
| --- | --- | --- |
| taxon_id | integer | ID of a *Taxon* (unique) |
| taxon_species | integer | ID of a *Taxon*, which was assigned as species level |
| taxon_genus | integer | ID of a *Taxon*, which was assigned as genus level |
| taxon_family | integer | ID of a *Taxon*, which was assigned as family level |

### TaxonReference table

Each *Taxon* might get a reference. Currently, this is used to track the origin of the *Taxon* name (i.e., which primary source was used first with this *Taxon*). Note that *Taxa*, generated from taxonomic classification (see **Methods**), are associated with “taxospace” reference. See **Table 26** for the description of individual columns.

**Table 26:** Column names and types for table *TaxonReference*.

| Column name | Data type | Description |
| --- | --- | --- |
| taxon_id | integer | ID of a *Taxon* |
| reference_id | integer | ID of a Reference |

**Table 27:** Column names and types for table *Traits*.

| Column name | Data type | Description |
| --- | --- | --- |
| trait_id | integer | ID of a *Trait* (unique) |
| trait_domain_id | integer | ID of a *Trait Domain* |
| trait_name | character | Name of the trait from the primary source. See **Methods** for the details about the specific columns used from primary sources. |

### *Traits* table

The *Traits* table contains the list of functional traits currently contained in VegVault. The table contains one *Trait* per row, with each *Trait* may contain: a unique *Trait* ID (“trait_id”), original *Trait name* from primary source (“trait_name”; see **Methods** for the details about the specific columns used from primary sources), and *Trait Domain* (“trait_domain_id”). See **Table 20** for the description of individual columns.

### TraitsDomain table

Traits are grouped into *Trait Domains* to allow for easier aggregation of *Traits* across data sources (see **Methods** section). In total, six *Trait Domains* are present: “Stem specific density”, “Leaf nitrogen content per unit mass”, “Diaspore mass”, “Plant height”, “Leaf area”, “Leaf mass per area”, following Diaz et al. (2016)^36^. See **Table 3** for an overview of *Traits* in various *Trait Domains* and the number of *Trait* values. Yet, it is up to the user to decide how to further aggregate trait values if multiple trait samples of one *Trait Domain* are available for the same environmental or taxonomic entity. See **Table 28** for the description of individual columns.

**Table 28:** Column names and types for table *TraitsDomain*.

| Column name | Data type | Description |
| --- | --- | --- |
| trait_domain_id | integer | ID of a *Trait Domain* (unique) |
| trait_domain_name | character | Name of the *Trait Domain* from Diaz et al. (2016)<sup>36</sup> |
| trait_domanin_description | character | Additional information about the *Trait Domain* |

### TraitsValue table

In general, data of functional traits of vegetation taxa follow the same structure of the *Dataset* and *Samples* obtained directly from the *Dataset Source-Types*. Therefore, the *TraitsValue* table contains not only the actual measured value of *Trait* observation but also information about linking information across *Datasets*, *Samples*, and *Taxa.* This comprehensive linkage ensures that each *Trait* value is accurately associated with its relevant ecological, environmental and taxonomic context. See **Table 29** for the description of individual columns.

**Table 29:** Column names and types for table *TraitsValue*.

| Column name | Data type | Description |
| --- | --- | --- |
| trait_id | integer | ID of a *Trait* |
| dataset_id | integer | ID of a *Dataset* |
| sample_id | integer | ID of a *Sample* |
| taxon_id | integer | ID of a *Taxon* |
| trait_value | numeric | Value of specific measured observation of *Trait*. See **Methods** for the details about the specific columns used from primary sources. |

### TraitsReference table

To ensure clarity and reproducibility, each *Trait* in the VegVault can have additional *References* beyond the general *Dataset* and *Sample References*. These *Trait*-specific *References* provide detailed provenance and citation information, supporting rigorous scientific research and enabling users to trace the origins and validation of each trait value. See **Table 30** for the description of individual columns.

**Table 30:** Column names and types for table *TraitsReference*.

| Column name | Data type | Description |
| --- | --- | --- |
| trait_id | integer | ID of a *Trait* |
| reference_id | integer | ID of a *Reference* |

### *AbioticVariable* table

As the VegVault contains abiotic variables from several primary sources, the *AbioticVariable* table contains descriptions of abiotic variables (“abiotic_variable_name*”*), their units (“abiotic_variable_unit”), and measurement details (“measure_details”). See **Table 4** for the overview of abiotic variables and their primary source, and **Table 31** for the description of individual columns.

**Table 31:** Column names and types for table *AbioticVariable*.

| Column name | Data type | Description |
| --- | --- | --- |
| abiotic_variable_id | integer | ID of a *Abiotic Variable* |
| abiotic_variable_name | character | Name of a *Abiotic Variable* from primary source |
| abiotic_variable_unit | character | Unit of a *Abiotic Variable* |
| measure_details | character | Additonal details about *Abiotic Variable* |

### AbioticData table

The table holds the actual values of abiotic variables (the units are the same for each *Abiotic Variable*). See **Table 32** for the description of individual columns.

**Table 32:** Column names and types for table *AbioticData*.

| Column name | Data type | Description |
| --- | --- | --- |
| sample_id | integer | ID of a *Sample* |
| abiotic_variable_id | integer | ID of an *Abiotic Variable* |
| abiotic_value | numeric | The actual value of abiotic data |

### AbioticVariableReference table

Each *Abiotic Variable* can have a separate *Reference*, in addition to a *Dataset* and *Sample*. See **Table 33** for the description of individual columns.

**Table 33:** Column names and types for table *AbioticVariableReference*.

| Column name | Data type | Description |
| --- | --- | --- |
| abiotic_variable_id | integer | ID of an *Abiotic Variable* |
| reference_id | integer | ID of a *Reference* |

### AbioticDataReference table

As abiotic data is stored in a special *Dataset Type*—*gridpoints*, we have also pre-calculated the spatio-temporal distance of *gridpoints* to other Datasets. See **Methods** for details about the creation of *gridpoints*. There, the *AbioticDataReference* table contains information about specific distances among *Samples* for both temporal (age in years) and geographic (distance in km). See **Table 34** for the description of individual columns.

**Table 34:** Column names and types for table *AbioticDataReference*.

| Column name | Data type | Description |
| --- | --- | --- |
| sample_id | integer | ID of non-*gridpoint* Sample |
| sample_ref_id | integer | ID of *gridpoint* Sample |
| distance_in_km | integer | Distance among samples expressed in kilometres |
| distance_in_years | integer | Distance among samples expressed in years |

**Table 35:**
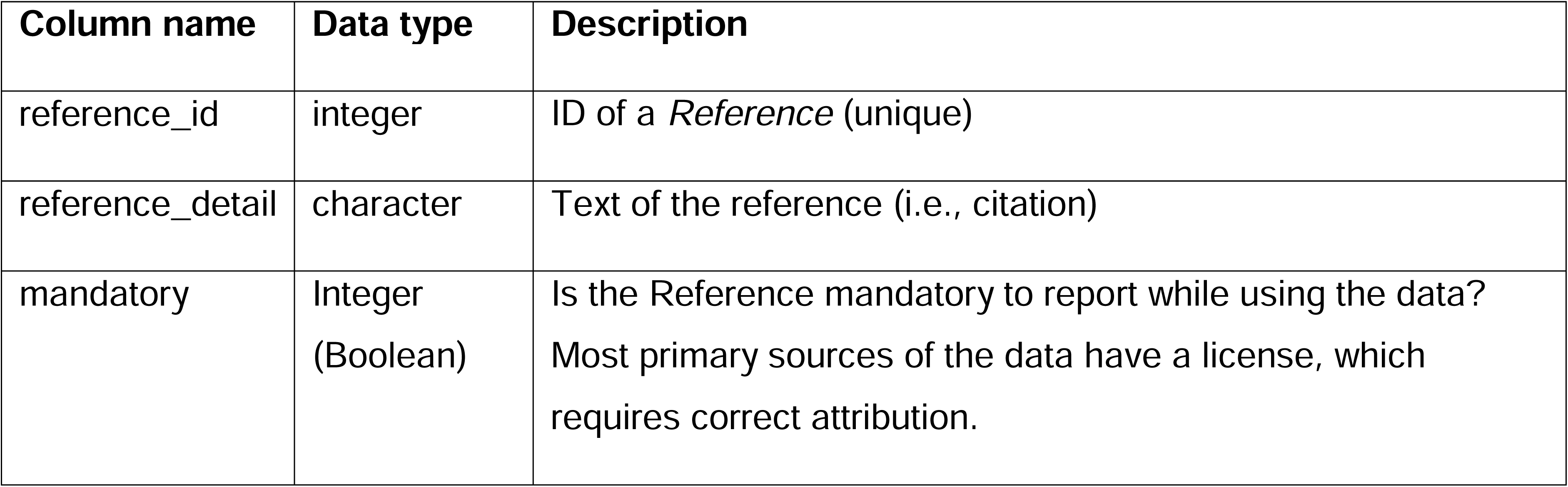
Column names and types for table *References*.

### *References* table

This table contains all *References*, independent of the source of the reference and the type of data. Each row contains a single *Reference* that is then linked to the type of data which is being referenced. This allows to have a single *Reference* to be used across data types, but also one data point having many different references.

Moreover, most primary sources of the data have a license, which requires correct attribution. Therefore, each *Reference* has information on whether such a *Reference* needs to be cited while using the specific data (“mandatory” column).

### Technical Validation

The data validity of VegVault is driven by the completeness, accuracy, and reliability of the source data. Several steps have been undertaken to ensure that VegVault provides high-quality data for research applications. For the fossil pollen records, strict data-quality rules have been applied by using the *FOSSILPOL* workflow^16^. The data preparation and execution are backed up by open-source publicly available code versioned and controlled by a git Tag system.

## Usage Notes

### Guidelines for data reuse and update

Due to the version system of VegVault, the dataset is expected to be updated using new data and/or changing the process of data migration and processing. Therefore, we recommend citing the latest VegVault version with the corresponding DOI, see <u>bit.ly/VegVault</u> for the latest release of the dataset (v1.0.0 can be found here: doi.org/10.48700/datst.t21cz-2nq07).

As the primary data sources are held under various licences, VegVault has a *CC BY 4.0* Licence (https://creativecommons.org/licenses/by/4.0/), allowing the copying and redistribution of the material as long as correct attribution is given. We recommend that users carefully review the restrictions associated with data usage. In addition, when citing the usage of VegVault, users should always cite the primary sources of the data, as well as any individual datasets. For those purposes, we created a function ‘*vaultkeepr::get_references()*’ within the {*vaultkeepr*} R package (see below) that will output a list of recommended references for the specific data compilation created by the user.

### Limitations and considerations for future research

Potential limitation lies in the completeness, accuracy, and reliability of the primary data sources. For example, it has been suggested that improper data practices erode the quality of global ecological databases such as *TRY*^47^, which have been used in VegVault. We do not attempt to solve these problems as VegVault’s main purpose is the re-usability of the data.

Next, as VegVault database contains more than 100GB of data, we advise the users to make sure that their machine has enough space on their drives, as downloads can be challenging on certain machines (e.g., a laptop).

Finally, to increase the accessibility of the data, we have also deposited the flat version of the database (i.e. individual tables saved as CSV files), accessible under DOI: <u>10.48700/datst.thktf-</u> <u>5rr35</u>. However, note that many of those files are larger than 1GB and some machines do not have enough memory (RAM) to fully load them.

### {vaultkeepr}: R-package for accessing the VegVault database

In order to make usage of the VegVault as easy as possible and facilitate access to SQLite databases, we have developed an R-package called {*vaultkeepr*}, providing a suite of functions to effectively interact with the database directly from the R programming environment^48^.

Functions include accessing the database, extracting datasets, filtering samples, and accessing specific taxa and traits. Therefore, we suggest all users to use the SQLite database accompanied by {vaultkeepr} R package to retrieve only project-specific data without the need to load data into memory.

This package is a well-tested interface (>95% code coverage). See **Figure 9** for details about individual steps to retrieve the data from VegVault. For more about the package, the function documentation and examples of code usage, please see {*vaultkeepr*} website (https://bit.ly/vaultkeepr). Here we present 3 examples of potential projects and how to obtain data for such projects from VegVault using {*vaultkeepr*} R-package. Note that we are specifically not trying to do any analysis, only presenting a way to obtain data, which can be used for such follow-up projects:

**Figure 9.**
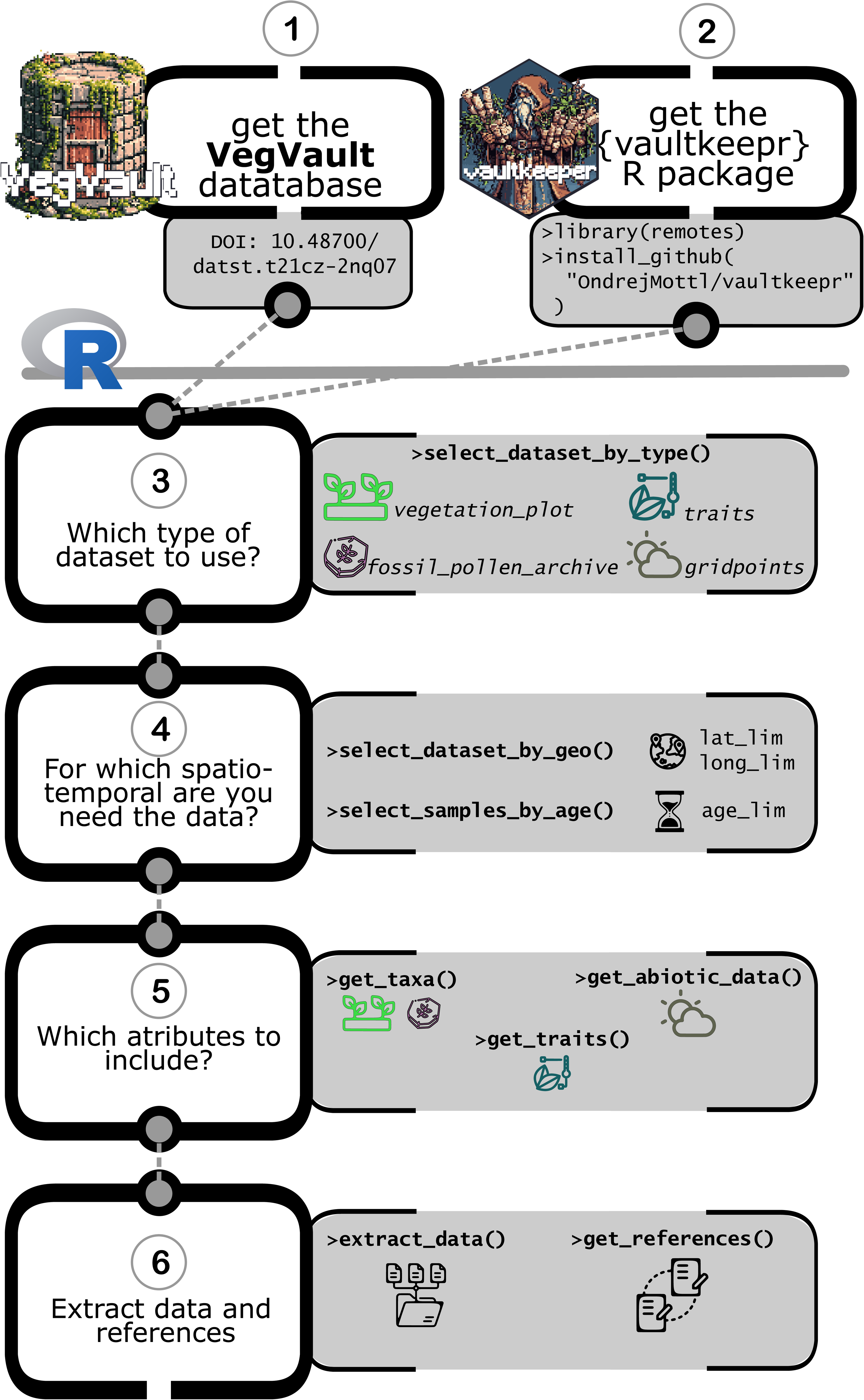
Schematic workflow of accessing and extracting data from VegVault database using {vaultkeepr} R package. **<u>1)</u>** Download VegVault database (v1.0.0; DOI: 10.48700/datst.t21cz-2nq07); **<u>2)</u>** Download {vaultkeepr} R package from GitHub (current version 0.0.6; DOI: 10.5281/zenodo.14964737); **<u>3)</u>** In the R programming environment^48^, the user has to select which type(s) of datasets they would like to extracted. A minimum of one *Dataset Type (see* **Data Records** section for more details) must be specified when using the ‘*vaultkeepr::select_dataset_by_type()*’ function; **<u>4)</u>** When the user is interested in data from a specific region, they can specific the geographical coordinates of a rectangular extent using the ‘*vaultkeepr::select_dataset_by_geo()*’ function to only get *Samples* from the individual *Datasets* that were recorded within the region of interest. Additionally, the user can specify the temporal focus by filtering data within a specific period using the ‘*vaultkeepr::select_samples_by_age()*’; **<u>5)</u>** The user may also specify any further attributes to be added to the data compilation. Specifically: **’***<u>get_taxa()</u>*’—When extracting contemporary or past vegetation (fossil pollen) records, the user most likely wishes to add information about the abundance of individual *Taxa* in each *Sample*. To do so, the user can use the ‘*vaultkeepr::get_taxa()*’ function. In addition, the user can standardise the taxonomy, so that the extracted *Taxa* can be compared. The parameter ‘*classify_to*’ within the ‘*vaultkeepr::get_taxa()*’ function allows the user to specify a taxonomic level (species, genus or family) on which the data should be comparable (see **Methods** about *Taxa* classification). Furthermore, the user can select specific *Taxa* based on taxonomy by using the ‘*vaultkeepr::select_taxa_by_name()*’ function. ‘*<u>get_traits()</u>*’—When wishing to link the *Trait* data with the vegetation records, the user can use ‘*vaultkeepr::get_traits()’* function to extract all *Trait Samples* of the earlier specified spatio-temporal extent. Moreover, similarly to the ‘*vaultkeepr::get_taxa()*’ function, the user can specify the taxonomic level to which the data should be standardised by using the ‘*classify_to*’ parameter. Please note that the user has to decide in their further analysis how they want to aggregate the measured traits per taxonomic group, e.g. by taking the mean. Further, the user can select a specific *Trait Domain* (of the six available; see **Methods** about details) by using the ‘*vaultkeepr::select_traits_by_domain_name()’* function. *’get_abiotic_data()’—*When wishing to link vegetation or trait records with abiotic data, the abiotic data can be obtained by the ‘*vaultkeepr::get_abiotic_data()’* function. The user can specify in which mode the abiotic data should be linked, which can be either “*nearest*” (i.e., geographically closest records), “*mean*” or “*median*” (summarising all abiotic records within a set geographical and/or temporal distance). This can be further tweaked by its parameters ‘*limit_by_distance_km*’ and ‘*limit_by_age_years*’. Furthermore, specific abiotic variables can be chosen by the ‘*vaultkeepr::select_abiotic_var_by_name()’* function; **<u>6)</u>** When defined all specifications for data extraction, the user can execute the extraction using ‘*vaultkeepr::extract_data()*’. This will result in a “ready-for-analyses” data compilation. Moreover, the user can use the ‘*vaultkeepr::get_references()*’ function to obtain all references required (and/or suggested) for such compilation (see **Internal Dataset Structure** section for more details about the hierarchical structure of references). Finally, see the **Usage Notes** section for examples of data processing and extracting using {vaultkeepr} R package.

**Figure 10A:** The first example demonstrates how to retrieve data for the genus *Picea* across North America by selecting both contemporary and fossil pollen plot datasets, filtering samples by geographic boundaries and temporal range (0 to 15,000 cal yr BP), and harmonizing taxa to the genus level. The resulting dataset allows users to study spatiotemporal patterns of *Picea* distribution over millennia.

**Figure 10B:** In the second example, the project aims to do species distribution modelling for plant taxa in the Czech Republic based on contemporary vegetation plot data and mean annual temperature. The project includes selecting datasets and extracting relevant abiotic data.

**Figure 10C:** The third example focuses on obtaining data to be able to reconstruct plant height for South and Central America between 6-12 cal kyr BP (thousand calibrated years before present). This example project showcases the integration of trait data with paleo-vegetation records to subsequently study historical vegetation dynamics and functional composition of plant communities.

**Figure 10.**
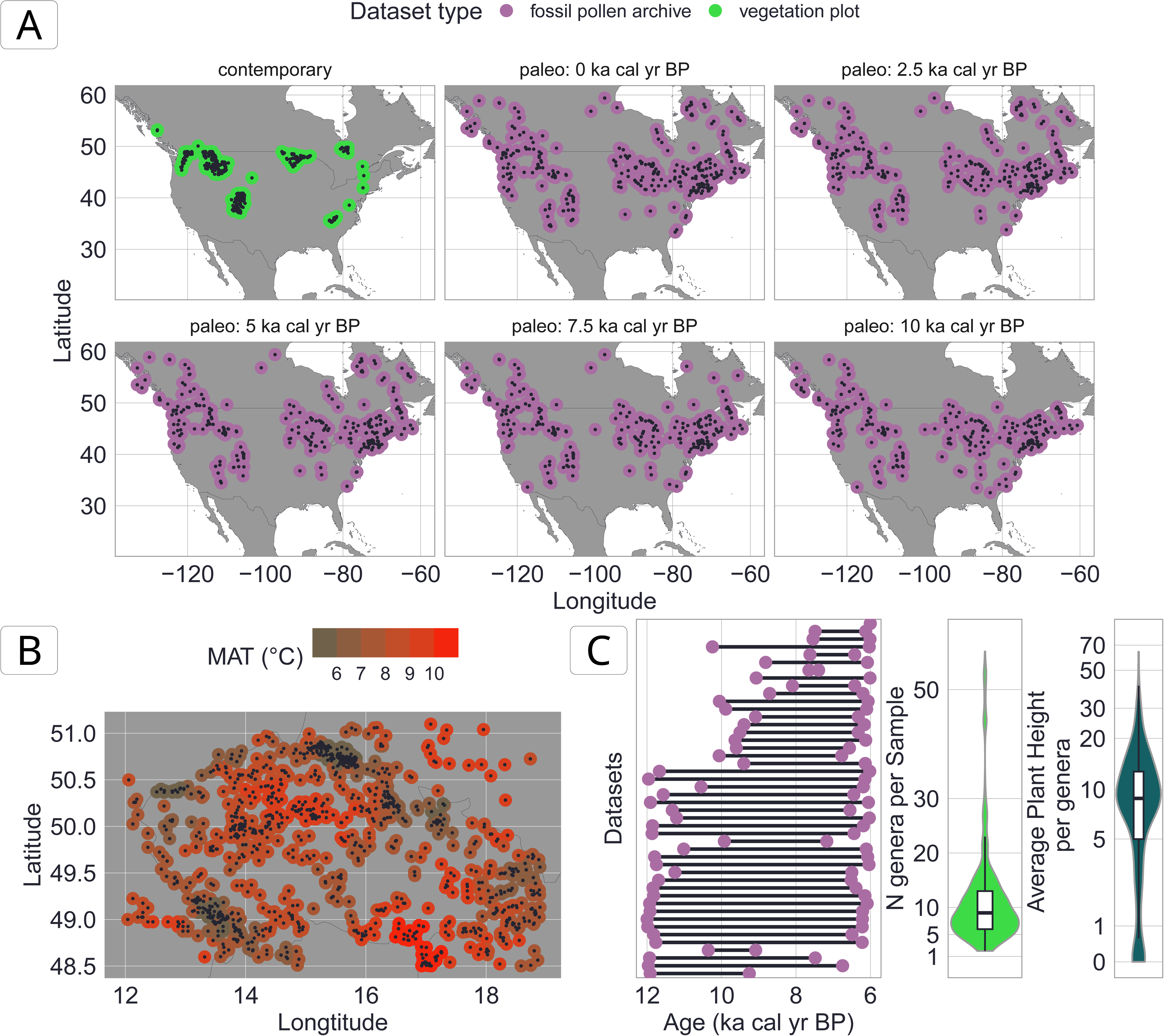
Visualisation of example data usage of VegVault using {*vaultkeepr*} package. See **Usage Notes** section for details about the aim of each data compilation. **A**: The individual points represent vegetation data (*Samples* for fossil pollen or contemporary vegetation plots) within each age time slice. **B**: Each point represents one contemporary vegetation plot. The colour represents the Mean Annual Temperature (MAT) associated with that plot. Note that several vegetation plots can have the same value due to sharing the same closest *gridpoint* (see **Figure 2**). **C**: The left figure represents the temporal length of fossil pollen *Datasets*, which are available for Central and South America between 6 and 12 ka BP (thousand years before present). The middle figure represents the distribution of various genera present in this data assembly. The right figure represents the distribution of average *Plant height Trait* per each genus present in the data assembly. The code to reproduce these figures can be accessed from the GitHub repository titled *VegVault* (https://github.com/OndrejMottl/VegVault/tree/v1.0.0-website_rewiev), which can be accessed as DOI: 10.5281/zenodo.16987345.

## Data Availability

The VegVault v1.0.0 relational database (SQLite; ∼110 GB) is publicly archived in the National Repository (‘REPO’) and available at https://doi.org/10.48700/datst.t21cz-2nq07. The deposit consists of a single SQLite file comprising 31 linked tables with 87 fields, containing >480,000 datasets, >13,000,000 samples, >100,000 taxa, six vegetation trait domains, >11,000,000 trait values, and eight abiotic variables. The database schema and field definitions are detailed in the manuscript (**Data Records**, Figure 4; **Table 5-35**). The database can be queried with any SQLite-compatible client or accessed programmatically via the companion RR package {vaultkeepr} (see **Usage Notes**). Version history and change logs are stored in the internal *version_control* table, and subsequent releases will be issued as versioned updates in the same repository.

## Code Availability

The VegVault database is built on SQLite, chosen for its robustness and compatibility with various data types. The VegVault v1.0.0 is available at the *National Repository*^44^ (‘*REPO*’) under DOI <u>10.48700/datst.t21cz-2nq07</u>. The database itself is under *CC B 4.0* licence, see **Usage Notes** section.

VegVault utilizes several tools and software to facilitate data management and access. All code used to generate the VegVault dataset is publicly available under the MIT licence. The GitHub repository *VegVault* used to integrate all data together and migrate them into SQLite database (can be accessed as DOI: <u>10.5281/zenodo.16987345</u>). See **Methods** section and Figure 1 for details about individual GitHub repositories.

To enable easy access and manipulation of data, {*vaultkeepr*} R package has been developed as an open access software under MIT licence (https://bit.ly/vaultkeepr).

The code to create the figures in this manuscript is available under DOI:10.5281/zenodo.16987345.

## Acknowledgements

OM is funded by the *Czech Science Foundation PIF* grant (GN23-06386I), by the *Charles University Research Centre program* (UNCE/24/SCI/006), and by the *Institutional Support for Science and Research* of the Ministry of Education, Youth and Sports of the Czech Republic. SGAF is funded by the *Trond Mohn Research Foundation* (TMF) and the University of Bergen by starting grant no. TMS2022STG03).

## Author contributions

OM conceived the study and programmed all related code. SGAF provided strategic guidance during the development of the project and manuscript, including decisions on data presentation and structure. FG, IS, and SGAF provided critical feedback on the technical infrastructure and database design. OM and SGAF wrote the first draft of the manuscript. SGAF, FG, and IS contributed to multiple rounds of revision. All authors approved the final version of the manuscript.

## Supplementary Tables

**Table S1:**
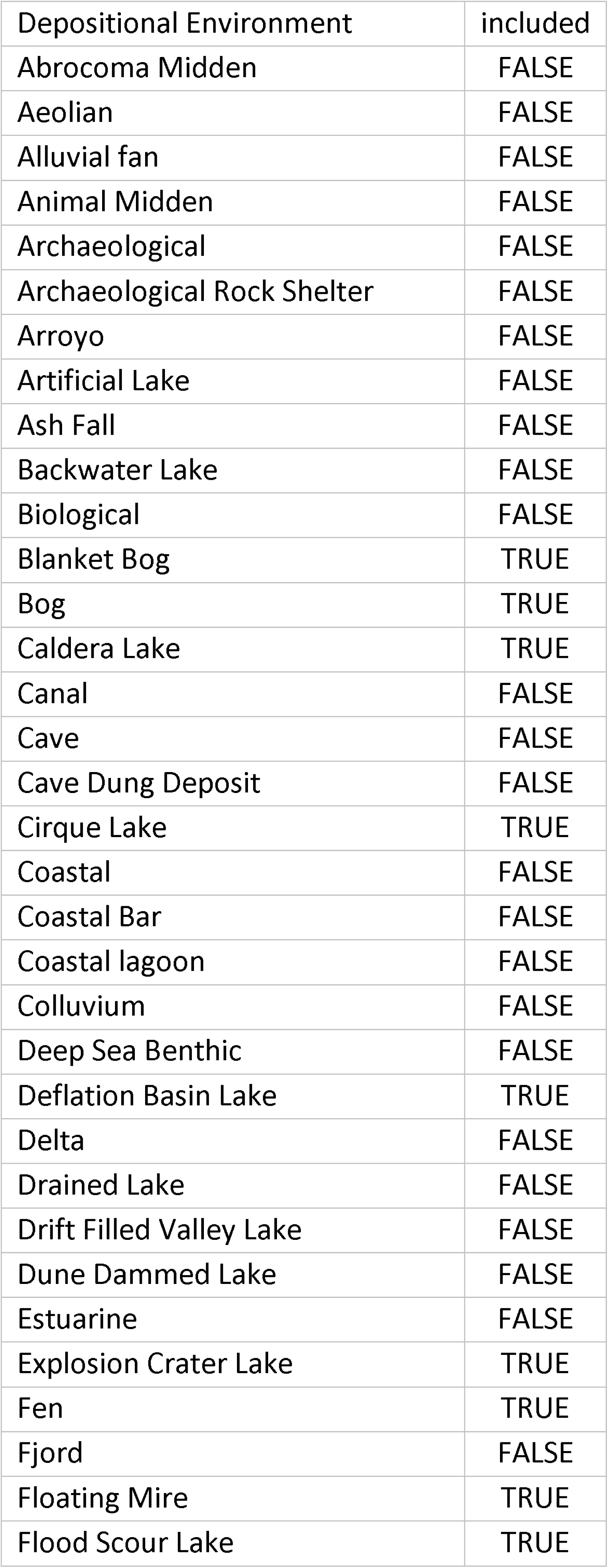
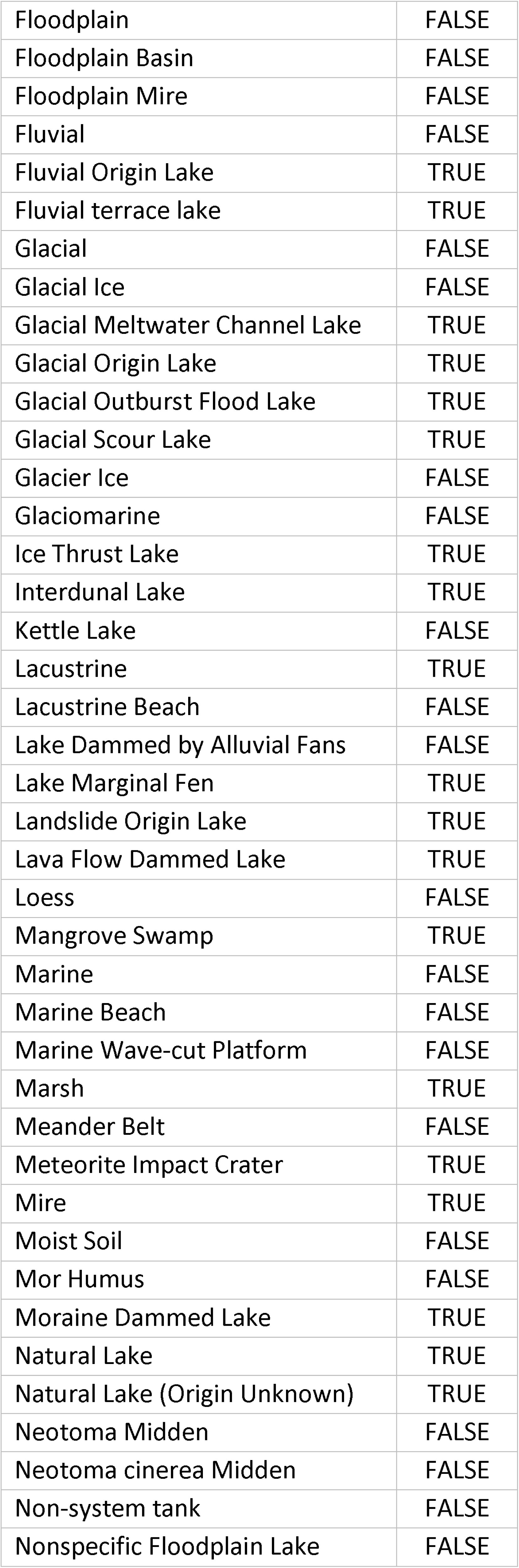
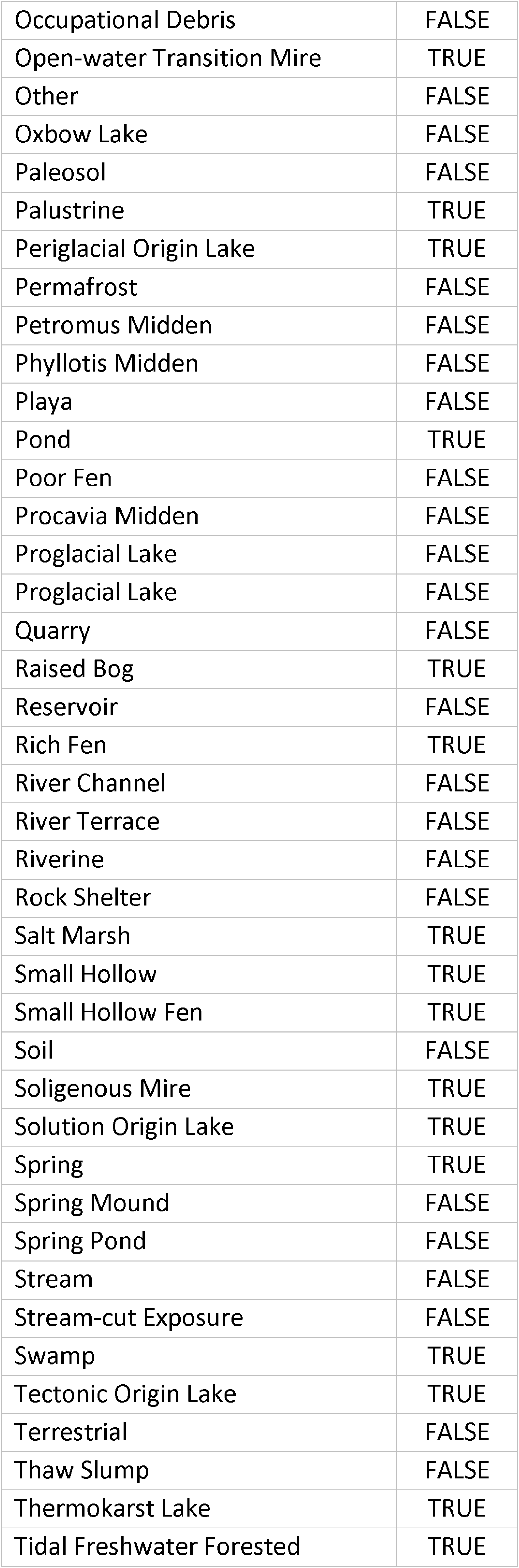
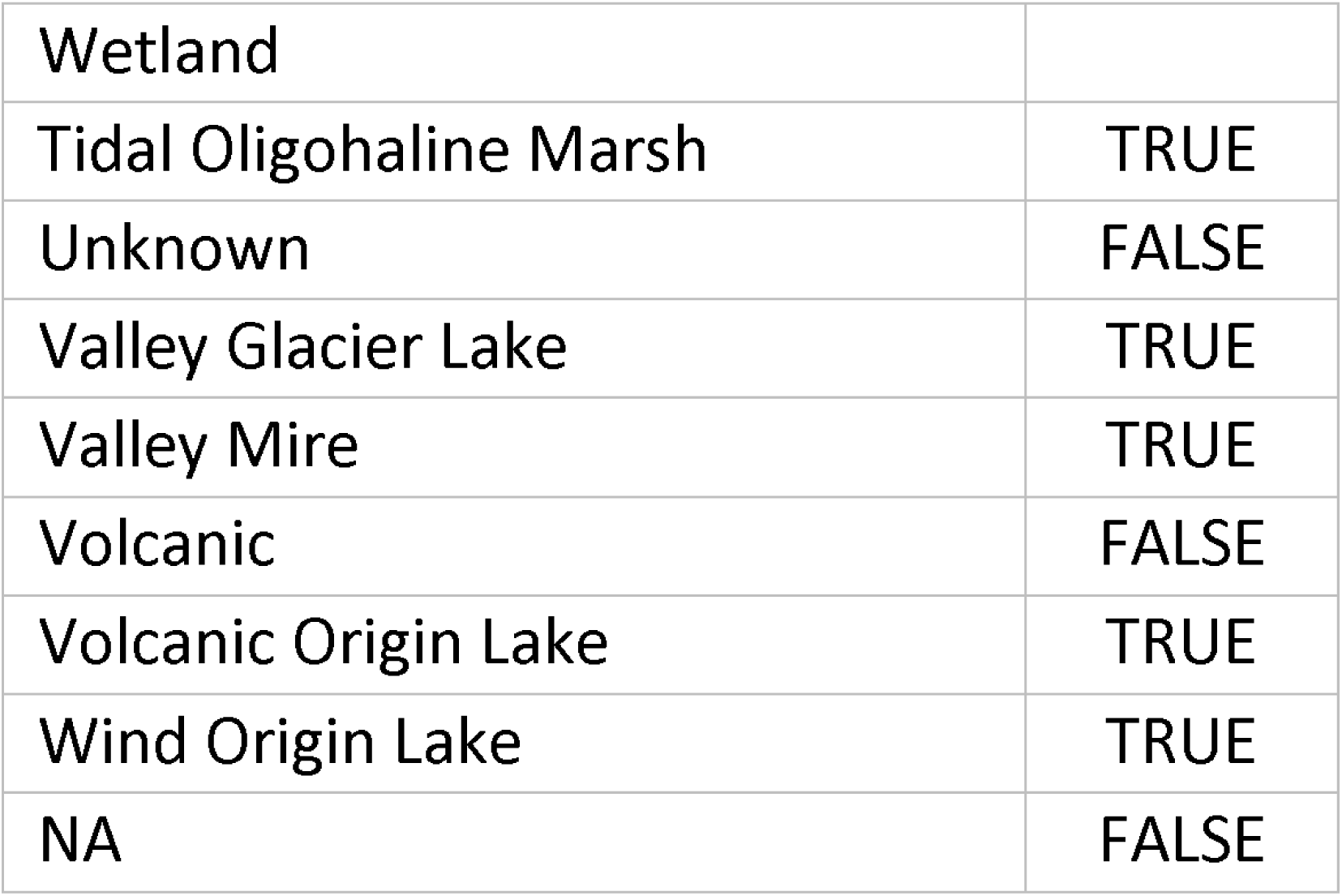
A list of selected depositional environments used during the processing of fossil pollen data.

**Table S2:** A list of selected ecological groups used during the processing of fossil pollen data.

| Ecological group | included |
| --- | --- |
| ACAR | FALSE |
| ACRI | FALSE |
| ALGA | FALSE |
| ANAC | FALSE |
| ANIM | FALSE |
| AQBR | FALSE |
| AQVP | FALSE |
| ARTH | FALSE |
| BIOM | FALSE |
| BRYZ | FALSE |
| CHAR | FALSE |
| CHRM | FALSE |
| CLAD | FALSE |
| COLE | FALSE |
| COPE | FALSE |
| DIAT | FALSE |
| DINO | FALSE |
| DIPT | FALSE |
| ELEM | FALSE |
| EMBR | FALSE |
| FLWO | FALSE |
| FORM | FALSE |
| FUNG | FALSE |
| GPOD | FALSE |
| INSE | FALSE |
| LABO | FALSE |
| LEPI | FALSE |
| MANG | TRUE |
| MNRL | FALSE |
| NEMA | FALSE |
| PALM | TRUE |
| PLNT | TRUE |
| PLOI | FALSE |
| PROT | FALSE |
| PSED | FALSE |
| PTER | FALSE |
| ROTI | FALSE |
| SEED | FALSE |
| SPNG | FALSE |
| SUCC | TRUE |
| TEAM | FALSE |
| TRSH | TRUE |
| UNID | FALSE |
| UPBR | FALSE |
| UPHE | FALSE |
| VACR | FALSE |
| VASC | TRUE |
| NA | FALSE |

**Table S3:** A list of valid control types used during the age-depth modelling process during the processing of fossil pollen data.

| Chronology control type | included | calibrated |
| --- | --- | --- |
| Ambrosia rise | FALSE | FALSE |
| Annual laminations (varves) | TRUE | FALSE |
| Annual laminations (varves)/Sedimentation rate | TRUE | FALSE |
| Argon-argon | FALSE | FALSE |
| Biostratigraphic | FALSE | FALSE |
| Biostratigraphic, pollen | FALSE | FALSE |
| Biostratigraphic, pollen, Allerød/Younger Dryas | FALSE | FALSE |
| Biostratigraphic, pollen, Older Dryas/Allerød | FALSE | FALSE |
| Biostratigraphic, pollen, Oldest Dryas/Bølling | FALSE | FALSE |
| Biostratigraphic, pollen, Pleniglacial/Meiendorf | FALSE | FALSE |
| Biostratigraphic, pollen, Younger Dryas/Holocene | FALSE | FALSE |
| Biostratigraphic, pollen, regional | FALSE | FALSE |
| Collection date | TRUE | FALSE |
| Core bottom | FALSE | FALSE |
| Core top | TRUE | FALSE |
| Core top, estimated | TRUE | FALSE |
| Cultural | FALSE | FALSE |
| Deglaciation | FALSE | FALSE |
| European settlement horizon | FALSE | FALSE |
| Event stratigraphic | FALSE | FALSE |
| Extrapolated | FALSE | FALSE |
| Geochronologic | FALSE | FALSE |
| Geologic time scale | FALSE | FALSE |
| Geologic time-scale boundary | FALSE | FALSE |
| Guess | TRUE | FALSE |
| Heeb & Welten (1972) pollen boundary | FALSE | FALSE |
| Heinrich stadials | FALSE | FALSE |
| Historical documentation | TRUE | FALSE |
| Historical fire | TRUE | FALSE |
| IDW-2d Picea decline | FALSE | FALSE |
| IDW-2d Ulmus decline | FALSE | FALSE |
| Infrared Stimulated Luminescence (IRSL) | FALSE | FALSE |
| Interpolated | FALSE | FALSE |
| Introduced Pinus rise | FALSE | FALSE |
| Isotopic excursion | FALSE | FALSE |
| Lead-210 | TRUE | FALSE |
| Marine isotope stage boundary | FALSE | FALSE |
| Marine isotope stage event | FALSE | FALSE |
| Marine isotope stages | FALSE | FALSE |
| Optically stimulated luminescence | TRUE | FALSE |
| Other absolute dating methods | FALSE | FALSE |
| Other dating methods | FALSE | FALSE |
| Oxygen-18 | TRUE | FALSE |
| Palaeomagnetic | TRUE | FALSE |
| Picea decline | FALSE | FALSE |
| Pre-EuroAmerican settlement horizon | FALSE | FALSE |
| Quaternary event classification | FALSE | FALSE |
| Radiocarbon | TRUE | TRUE |
| Radiocarbon, average of two or more dates | TRUE | TRUE |
| Radiocarbon, calibrated | TRUE | FALSE |
| Radiocarbon, calibrated from calendar years | TRUE | FALSE |
| Radiocarbon, calibrated, Bayesian modelled | TRUE | FALSE |
| Radiocarbon, infinite | TRUE | FALSE |
| Radiocarbon, reservoir correction | TRUE | TRUE |
| Salsola rise | FALSE | FALSE |
| Section top | TRUE | FALSE |
| Sediment stratigraphic | FALSE | FALSE |
| Stratigraphic | FALSE | FALSE |
| Tephra | TRUE | FALSE |
| Tsuga decline | FALSE | FALSE |
| Uranium-series | TRUE | FALSE |
| W.O. van der Knaap pollen boundary | FALSE | FALSE |
| Welten (1982) pollen boundary | FALSE | FALSE |

## References

1. Fordham, D. A. et al. Using paleo-archives to safeguard biodiversity under climate change. Science 369, eabc5654 (2020) 10.1126/science.abc5654.

2. Nieto-Lugilde, D. et al. Time to better integrate paleoecological research infrastructures with neoecology to improve understanding of biodiversity long-term dynamics and to inform future conservation. Environ. Res. Lett. 16, 095005 (2021) 10.1088/1748-9326/ac1b59.

3. Reitalu, T., Kuneš, P. & Giesecke, T. Closing the gap between plant ecology and Quaternary palaeoecology. Journal of Vegetation Science 25, 1188–1194 (2014) 10.1111/jvs.12187.

4. GBIF.org. GBIF Home Page. (2024) https://www.gbif.org.

5. Bruelheide, H. et al. sPlot – A new tool for global vegetation analyses. Journal of Vegetation Science 30, 161–186 (2019) 10.1111/jvs.12710.

6. Botanical Information and Ecology Network (BIEN). BIEN: Botanical Information and Ecology Network. http://bien.nceas.ucsb.edu/bien/ (n.d.).

7. Williams, J. W. et al. The Neotoma Paleoecology Database, a multiproxy, international, community-curated data resource. Quat. res. 89, 156–177 (2018) 10.1017/qua.2017.105.

8. Kattge, J. et al. TRY plant trait database – enhanced coverage and open access. Global Change Biology 26, 119–188 (2020) 10.1111/gcb.14904.

9. Karger, D. N. et al. Climatologies at high resolution for the earth’s land surface areas. Sci Data 4, 170122 (2017) 10.1038/sdata.2017.122.

10. Fick, S. E. & Hijmans, R. J. WorldClim 2: new 1-km spatial resolution climate surfaces for global land areas. International Journal of Climatology 37, 4302–4315 (2017) 10.1002/joc.5086.

11. Schipper, E. L. F., Dubash, N. K. & Mulugetta, Y. Climate change research and the search for solutions: rethinking interdisciplinarity. Clim Change 168, 18 (2021) 10.1007/s10584-021-03237-3.

12. Intergovernmental Panel On Climate Change. Climate Change and Land: IPCC Special Report on Climate Change, Desertification, Land Degradation, Sustainable Land Management, Food Security, and Greenhouse Gas Fluxes in Terrestrial Ecosystems. (Cambridge University Press, 2022). 10.1017/9781009157988 https://www.cambridge.org/core/product/identifier/9781009157988/type/book.

13. Smith, J. et al. BioDeepTime: A database of biodiversity time series for modern and fossil assemblages. Global Ecology and Biogeography 32, 1680–1689 (2023) 10.1111/geb.13735.

14. Weigelt, P., König, C. & Kreft, H. GIFT – A Global Inventory of Floras and Traits for macroecology and biogeography. Journal of Biogeography 47, 16–43 (2020) 10.1111/jbi.13623.

15. Sabatini, F. M. et al. sPlotOpen – An environmentally balanced, open-access, global dataset of vegetation plots. Global Ecology and Biogeography 30, 1740–1764 (2021) 10.1111/geb.13346.

16. Flantua, S. G. A. et al. A guide to the processing and standardization of global palaeoecological data for large-scale syntheses using fossil pollen. Global Ecology and Biogeography 32, 1377–1394 (2023) 10.1111/geb.13693.

17. Batjes, N. H., Calisto, L. & de Sousa, L. M. Providing quality-assessed and standardised soil data to support global mapping and modelling (WoSIS snapshot 2023). Earth System Science Data 16, 4735–4765 (2024) 10.5194/essd-16-4735-2024.

18. Karger, D. N., Nobis, M. P., Normand, S., Graham, C. H. & Zimmermann, N. E. CHELSA-TraCE21k: Downscaled transient temperature and precipitation data since the last glacial maximum. (2020) 10.16904/envidat.211 https://www.envidat.ch/dataset/chelsa_trace.

19. Karger, D. N., Nobis, M. P., Normand, S., Graham, C. H. & Zimmermann, N. E. CHELSA-TraCE21k – high-resolution (1km) downscaled transient temperature and precipitation data since the Last Glacial Maximum. Climate of the Past 19, 439–456 (2023) 10.5194/cp-19-439-2023.

20. GBIF Secretariat. GBIF Backbone Taxonomy. (2023).

21. Mottl, O. taxospace - v0.0.0.9003. 10.5281/zenodo.14699491 (2024) https://github.com/OndrejMottl/taxospace.

22. Mottl, O. vaultkeepr - v0.0.6. 10.5281/zenodo.14964737 (2025) https://github.com/OndrejMottl/vaultkeepr.

23. Mottl, O. et al. Global acceleration in rates of vegetation change over the past 18,000 years. Science 372, 860–864 (2021) 10.1126/science.abg1685.

24. Šímová, I., Ordonez, A. & Storch, D. The dynamics of the diversity–energy relationship during the last 21,000 years. Global Ecology and Biogeography 32, 707–718 (2023) 10.1111/geb.13649.

25. Brown, K. A. et al. Trait-based approaches as ecological time machines: Developing tools for reconstructing long-term variation in ecosystems. Functional Ecology 37, 2552–2569 (2023) 10.1111/1365-2435.14415.

26. Li, J. & Prentice, I. C. Global patterns of plant functional traits and their relationships to climate. Commun Biol 7, 1–14 (2024) 10.1038/s42003-024-06777-3.

27. Shipley, B., Dilkina, B. & McGuire, J. MegaSDM: Modelling Species Ranges in The Past And Future. Bulletin of the Florida Museum of Natural History 60, 115–115 (2023) 10.58782/flmnh.zwwl8127.

28. Pollock, L. J. et al. Understanding co-occurrence by modelling species simultaneously with a Joint Species Distribution Model (JSDM). Methods in Ecology and Evolution 5, 397–406 (2014) 10.1111/2041-210X.12180.

29. Ordonez, A. & Gill, J. L. Unravelling the functional and phylogenetic dimensions of novel ecosystem assemblages. Philosophical Transactions of the Royal Society B: Biological Sciences 379, 20230324 (2024) 10.1098/rstb.2023.0324.

30. Wieczynski, D. J. et al. Climate shapes and shifts functional biodiversity in forests worldwide. Proceedings of the National Academy of Sciences 116, 587–592 (2019) 10.1073/pnas.1813723116.

31. Zurell, D., Fritz, S. A., Rönnfeldt, A. & Steinbauer, M. J. Predicting extinctions with species distribution models. Cambridge Prisms: Extinction 1, e8 (2023) 10.1017/ext.2023.5.

32. Chacon, S. & Straub, B. Pro Git. (Springer Nature, 2014).

33. Haslett, J. & Parnell, A. A Simple Monotone Process with Application to Radiocarbon-Dated Depth Chronologies. Journal of the Royal Statistical Society Series C: Applied Statistics 57, 399–418 (2008) 10.1111/j.1467-9876.2008.00623.x.

34. Maitner, B. S. et al. The bien r package: A tool to access the Botanical Information and Ecology Network (BIEN) database. Methods in Ecology and Evolution 9, 373–379 (2018) 10.1111/2041-210X.12861.

35. Sabatini, F. M., Lenoir, J., Bruelheide, H. & sPlot Consortium. sPlotOpen – An environmentally-balanced, open-access, global dataset of vegetation plots. iDiv Data Repository (2021) 10.25829/idiv.3474-bb7k72.

36. Díaz, S. et al. The global spectrum of plant form and function. Nature 529, 167–171 (2016) 10.1038/nature16489.

37. Lam, O. H. Y. et al. ‘rtry’: An R package to support plant trait data preprocessing. Ecology and Evolution 14, e11292 (2024) 10.1002/ece3.11292.

38. Jentsch, helge, Weidinger, J. & Bobrowski, M. ClimDatDownloadR: Downloads Climate Data from Chelsa and WorldClim. (2025) 10.5281/zenodo.7924343 https://github.com/HelgeJentsch/ClimDatDownloadR.

39. Hijmans, R. J. Terra: Spatial Data Analysis. (2025). https://rspatial.org/.

40. Thuiller, W. On the importance of edaphic variables to predict plant species distributions – limits and prospects. Journal of Vegetation Science 24, 591–592 (2013) 10.1111/jvs.12076.

41. Zuquim, G., Costa, F. R. C., Tuomisto, H., Moulatlet, G. M. & Figueiredo, F. O. G. The importance of soils in predicting the future of plant habitat suitability in a tropical forest. Plant Soil 450, 151–170 (2020) 10.1007/s11104-018-03915-9.

42. Velazco, S. J. E., Galvão, F., Villalobos, F. & Júnior, P. D. M. Using worldwide edaphic data to model plant species niches: An assessment at a continental extent. PLOS ONE 12, e0186025 (2017) 10.1371/journal.pone.0186025.

43. Global Names Architecture. Global Names Resolver. https://resolver.globalnames.org/

44. CESNET. National Repository. https://data.narodni-repozitar.cz/ (2024).

45. Dillon, E. M. et al. Challenges and directions in analytical paleobiology. Paleobiology 49, 377–393 (2023) 10.1017/pab.2023.3.

46. Keil, P. et al. Patterns of beta diversity in Europe: the role of climate, land cover and distance across scales. Journal of Biogeography 39, 1473–1486 (2012) 10.1111/j.1365-2699.2012.02701.x.

47. Augustine, S. P., Bailey-Marren, I., Charton, K. T., Kiel, N. G. & Peyton, M. S. Improper data practices erode the quality of global ecological databases and impede the progress of ecological research. Global Change Biology 30, e17116 (2024) 10.1111/gcb.17116.

48. R Core Team. R: A Language and Environment for Statistical Computing. (R Foundation for Statistical Computing, Vienna, Austria, 2024). https://www.R-project.org/.

